# Why the adventitious roots of poplar are so colorful: RNAseq and metabolomic analysis reveal flavonols, flavones, and anthocyanins accumulation in canker pathogens-induced adventitious roots in poplar

**DOI:** 10.1101/2024.03.09.584208

**Authors:** Li Min, Fu Yuchen, Li Jinxin, Shen Wanna, Wang Li, Li Zheng, Zhang Shiqi, Liu Huixiang, Su Xiaohua, Zhao Jiaping

## Abstract

Recently, we observed a novel allometry on poplar stems, with copious colorful adventitious roots (ARs) induced by fungal canker pathogens. Here, we reveal chemical, physiological, and molecular mechanisms of AR coloration in a poplar-pathogen (*Valsa sordida*/*Botrosphaeria dothidea*) interaction system using our phloem girdling-inoculation system. Light-induced coloration in ARs: red/rosy under sunlight and milky white under shading. Chemical and metabolomic analyses indicated that numerous (93 in all 110) and high relative intensities/contents of flavonoids metabolites (mainly including flavonols, flavones, and anthocyanins class) accumulate in red ARs, some flavones and anthocyanins metabolites all contribute to the color of poplar ARs, and cyanidin-3-O-glucoside is the most abundant colorant. Integrated analysis of metabolomic and transcriptomic analysis suggested that sunlight exposure redirected metabolomic flux from the flavonoid biosynthesis pathway to the flavonols and flavones branch pathways, induced by the upregulation of FLS (flavonol synthase/flavanone 3-hydroxylase) and other structural genes. The anthocyanins metabolomic analysis and the downregulation of the ANS (anthocyanin synthase) gene illustrated a retard of metabolomic flux from leucoanthocyanidins to anthocyanidins; meanwhile, metabolomic results and the upregulation of gene BZ1 (Bronze 1, anthocyanin 3-O-glucosyltransferase) illustrated that sunlight triggered a rapid biosynthesis of anthocyanin metabolites in poplar ARs, which based on the substrate level of anthocyanidins. Transcriptomic and RT-qPCR analyses showed that transcriptional factor MYB113, HY5 (ELONGATED HYPOCOTYL5), and COP1 (Ring-finger protein CONSTITUTIVE PHOTOMORPHOGENIC1) genes positively regulate the expression of the flavonoid/anthocyanin biosynthesis structural genes (such as genes encoding BZ1, FLS, LAR, etc.) in both sunlight-exposed red ARs and white ARs after light exposure, suggesting sunlight induces anthocyanins biosynthesis through the interaction between “MBW” complex and COP1-HY5 module. Moreover, results also showed that 1 SPL gene (squamosa promoter-binding-like protein gene, target of miR156), one component of miR156-SPL module, downregulated in sunlight-exposed poplar ARs, implying the biosynthesis flavonoid/anthocyanin be regulated at the posttranscriptional level. Additionally, this study provides a potential AR experimental system for research on flavonoid/anthocyanin biosynthesis in tree species.

## 1. Introduction

Nature comprises diverse or even endless kinds/species of plants, animals, fungi, algae, and minerals, which are beautiful and colorful. For example, as the most common and abundant color originating from living organisms, almost all plant leaves are green, and the diverse and varied colors and shapes of flowers are associated with peace and beauty. The colors of plants can be divided into two categories: structural color and chemical color according to the principle of color rendering; and chemical color, also known as pigment color, is the color presented by organisms in nature through pigment production (Kinoshita and Yoshioka 2005; Kinoshita et al. 2002). For example, the content of chlorophyll in typical green leaves is generally higher than that of carotenoids; chlorophyll absorbs red and blue light of the spectrum and reflects only green light, giving leaves a greenish appearance (Glover and Whitney 2010). Plants produce a variety of pigment compounds (Fiehn 2002) that contribute to colors other than green, and pigmentation is mainly caused by the accumulation of flavonoids, anthocyanins, carotenoids, and betacyanins within plant epidermal cells (Zhao et al. 2012).

Flavonoids are composed of many phenylpropanoid secondary metabolites of plants, and they are biologically active and perform diverse functions in plants such as defense against biotic and abiotic stresses (Kozłowska and Szostak-Wegierek 2014; Nakabayashi et al. 2014). Flavonoids can be divided into the following classes according to their chemical structure: flavones, flavonols, anthocyanins, chalcones, isoflavones, flavanols, and flavanones (Shen et al. 2022). Flavonoids determine the color of plants involving a very wide range of colors, ranging from yellowish to blue. For example, flavonoids and isoflavones are responsible for the dark yellow color of red Chinese pear fruits (Ni et al. 2020), flavonoids, flavonol, and isoflavones participate in color synthesis in colored radishes (Zhang et al. 2020), whereas anthocyanins provide a wide variety of colors in most fruits and vegetables, ranging from bright red-orange to blue-violet colors (Cho et al. 2016; Lightbourn et al. 2007; Sass-Kiss et al. 2005; Tanaka et al. 2008).

The coloration of plants is a complex but finely tuned physiological and biochemical process. Studies indicate that biosynthesis and accumulation of secondary metabolites or pigments in plants are influenced by multiple environmental factors, such as light, temperature, nutritional status, water, soil type, microbial interactions, wounds, and phytohormones (Brunetti et al. 2019; Dixon and Paiva 1995; Fini et al. 2012; Fogelman et al. 2015; Kusano et al. 2011); however, light plays crucial roles in this process (Zoratti et al. 2014a; Zoratti et al. 2014b). Light quality affects flavonoid production and related gene expression in *Cyclocarya paliurus*: total leaf flavonoid contents are significantly higher under blue-, red-, and green light than under white light (Liu et al. 2018). Moreover, the production of flavonoids (anthocyanins and flavones) in *Silene littorea* is more plastic in photosynthetic tissues than in petals, and anthocyanin biosynthesis is more plastic than flavone production (Valle et al. 2018). For example, light induces accumulation of the anthocyanins by altering the expression of genes or transcript factors (TFs) involved in the production of anthocyanins, such as 4-coumarate coenzyme A ligase (4CL) and chalcone synthase (CHS) (Fang et al. 2020; Hong et al. 2016). Studies have illustrated that light induction of anthocyanins and flavonoids is regulated by R2R3-MYB TFs in red apples (Takos et al. 2006), *Petunia* (Albert et al. 2009), *Medicago truncatula* (Meng et al. 2019), and *Populus* (Kim et al. 2021; Peng et al. 2023), however, MYB10 is negatively related to the anthocyanins biosynthesis but promote secondary cell wall thicken in *Populus* (Jiang et al. 2022), and by *Pp*HYH in *Prunus persica* (Zhao et al. 2021).

In addition, the color of the newly sprouting roots of most plants is white, which might be the reason for the lack of pigment production or induction by light underground. For example, the hairy roots of *Echinacea purpurea* are yellow-white under dark conditions but light-grown hairy roots accumulated increased levels of anthocyanins, which became visible in the outer cell layer of the cortex as a ring of purple color (Abbasi et al. 2007). One research illustrated mechanical wounding induces flavonoids/anthocyanins production, for example, in transgenic poplar, overexpression of PdMYB118 promoted anthocyanin accumulation upon wound induction (Wang et al. 2020a). The biosynthesis and modification of flavonoids are the main and long-term strategies of *Dracaena cochinchinensis* response to wound stress (Liu et al. 2022a), red/blue light exposure gives a superior capacity wound-stress tolerance to *Arabidopsis thaliana* (by stimulating the synthesis of anthocyanin) (Mirzahosseini et al. 2020), and the increase of flavonoid, phenylpropanoid, sesquiterpenoid, and triterpenoid biosynthesis pathways might play important roles in wound-induced agarwood formation (Xu et al. 2023). However, the flavonoid/anthocyanin biosynthesis upon wound stress seems to be related to light induction or in natural conditions.

Poplar is one of the three major afforestation tree species, and the fungal stem canker diseases caused by *Botryosphaeria dothidea* and *Valsa sordida* are the most important forestry diseases in China. To investigate the physiology of poplar canker diseases, a phloem girdling-pathogen inoculation method was developed in our research (Xing et al. 2020). After 14-20 days, a novel allometric phenomenon, copious adventitious roots (ARs) with red color which sprouting from the inoculation sites of poplar stems, was observed in our research, aside from the common canker disease symptoms (such as bark lesions and canopy dieback), physiological and molecular performances (gas-exchange parameters, chlorophyll fluorescence parameters, carbohydrate contents, hydraulics, gene regulation, etc.) (Li et al. 2021; Li et al. 2019; Xing et al. 2020; Xing et al. 2022). To our best knowledge, there has been no report so far to find fungal pathogens-induced AR formation in plants. In light of the crucial roles of root development in the lifecycle of plants, the mechanisms of AR formation, and pathological and physiological effects of poplar ARs induced by canker pathogens have been investigated in our laboratory. Results showed that ARs produced in diverse poplar species and were induced by different fungi species, the accumulation of carbohydrates above the girdling site is necessary and the exogenous hormones (such as auxins) play crucial roles in the formation of poplar ARs (unpublished data); however, aimed at the understanding of coloration in poplar ARs, the mechanisms of flavonoid/anthocyanins biosynthesis are investigated and reported advanced.

Here, the morphological characteristics of the novel allometric growth of roots and AR formation induced by canker pathogens are first reported. Moreover, to reveal the physiology of AR coloration, the spectral features and pigment content (flavonoids, anthocyanins, and carotenoids), the metabolome of flavonoids and anthocyanins, transcriptome analysis of ARs induced by canker pathogens in sunlight or shaded conditions, and light-inducing pigment formation in shaded ARs were investigated in *Populus alba* var. *pyramidalis* saplings. Such study of poplar AR coloration will not only benefit the understanding biosynthesis of flavonoids/anthocyanins, but also provide interesting knowledge of the distribution, biosynthesis and accumulation of plant pigments, and insight into the formation mechanisms of pathogen-induced ARs in poplar.

## 2. Materials and methods

### 2.1 Plant materials, fungal pathogens, and AR formation

This study was carried out on 6-month-old *P. alba* var. *pyramidalis* saplings cultivated in pots containing a growth substrate of peat and perlite (v: v = 4:1) in the experimental field of the Chinese Academy of Forestry (Beijing, China). Some inoculation experiments were also carried out on branches of 4-year-old *P. alba* var. *pyramidalis.* The poplar saplings used in this study were healthy, without diseases or pests, and well-watered throughout the experiments. The poplar canker pathogen *B. dothidea* strain CZA and *V. sordida* strain CZC (Xing et al. 2020) were grown on 2.0% potato dextrose agar medium (2% potato extract, 2% dextrose, and 1.5% agar; pH 6.0) at 30 in the dark for 7 days.

#### 2.1.1 AR formation in sunlight and shade conditions

The medium with or without fungal mycelium was inoculated onto the stems of poplar saplings as described by Xing et al. (Xing et al. 2020). In brief, the stems of poplar saplings were girdled, and 1.2 cm samples of phloem and partial cambium were removed at 30 cm height above the growing substrate. Then, the media were cut into strips (1.2 cm in width and 2.5-3.0 cm in length) and inoculated at the girdle region. After pathogen inoculation, the girdling sites were wrapped with polyethylene (PE) or Parafilm^®^ film to maintain moisture. In shading experiments, the girdling sites were first wrapped with PE film and then covered fully with aluminum foil to prevent light-induced biosynthesis of color pigments. Three shading experiments were carried out in 2019-2022, and the saplings were inoculated with *V. sordida* (Vso) and *B. dothidea* (Bdo). In the present study, forty poplar saplings were inoculated, 20 saplings were wrapped using aluminum foil (S-Vso or S-Bdo), and other saplings without shading were used as control samples (Vso or Bdo).

#### 2.1.2 Light-induced color formation on shaded poplar ARs

In addition, the aluminum foil was removed from the six shaded poplar samples after the AR formation (25-30 days after inoculation, with white or light-colored ARs) and then exposed to sunlight. Morphological characteristics and color changes were observed for 3 days. The poplar ARs that formed under shaded conditions and the AR samples exposed to sunlight for 0, 1, and 3 days were collected for RNA extraction for RT-qPCR.

### 2.2 Morphology observation of pathogen-induced ARs in poplar

From 10 days after inoculation (DAI), the morphological features of ARs and their primordium (including the number of fibrous roots, the length and diameters of every fibril of ARs) around the girdling region of the poplar stem and the pathogenesis of canker disease were observed. To reveal the pigment distribution in pathogen-induced ARs, the microscopic characteristics of ARs (with or without shading) were observed. In this study, AR samples induced by both the pathogen *V. sordida* (S-Vso and Vso, at 25 DAI) and *B. dothidea* (S-Bdo and Bdo, at 25 DAI) were used. The AR slices were 50 μm in thickness and were not stained. Moreover, protoplasts of ARs (Vso) were prepared using 1.5% (wt/vol) Cellulase Onozuka R-10 (Yakult Pharmaceutical Ind. Co., Japan), 0.75% (wt/vol) Macerozyme Onozuka R-10 (Yakult Pharmaceutical Ind. Co., Japan) and 5% cell-wall degrading enzyme mix Viscozyme (Sigma-Aldrich, America).

### 2.3 Metabolite analyses

#### 2.3.1 Determination of contents of total flavonoids, anthocyanins, carotenoids, and proanthocyanidins

Total flavonoid, proanthocyanidin, and carotenoid contents were measured with plant flavonoid, proanthocyanidin, and carotenoid content assay kits (Solarbio, Beijing, China) according to the manufacturer’s protocol. In brief, approximately 0.1 g poplar shaded ARs (S-Vso) or poplar sunlight-exposed ARs (Vso) were ground to a powder for extraction of these three metabolites, and then their contents (mg/g) were measured at 470, 500, and 440 nm. In this study, all the poplar ARs samples were used in pigment content determination, metabolomic assays, and gene expression analysis were collected in dark conditions.

Approximately 0.1 g AR samples were extracted and measured at 530, 620, and 650 nm. The total anthocyanin content of ARs was determined by the pH-differential method (Giusti and Wrolstad 2001) as the following formula: 1) ODλ= (OD_530_ - OD_620_) - 0.1 (OD_650_ -OD_620_), 2) anthocyanin content (nmol/L) = 1000000 × OD ×ε^-1^× V × m^-1^, where OD is the optical density of anthocyanins at 530 nm, ε is the molar extinction coefficient of anthocyanin 4.62 × 10^6^, V is the total volume of the extract (mL), and m is the mass of the sample (g). In this study, the contents of total flavonoids, anthocyanins, proanthocyanidins, and carotenoids were evaluated in two independent experiments, with three independent biological replicates each.

#### 2.3.2 Metabolomic assays of flavonoids and anthocyanins

The poplar ARs induced by pathogen *V. sordida* inoculation in sunlight (Vso) and shade conditions (S-Vso) were used for flavonoids and anthocyanins metabolomic analysis. Three Vso samples and three S-Vso samples were analyzed in this study. Sample preparation, extraction analysis, metabolite identification, and quantification were performed by Wuhan MetWare Biotechnology Co., Ltd. (http://www.metware.cn/) following their standard procedures and previously fully described (Chen et al. 2013; Huang et al. 2021).

Flavonoid metabolic class was analyzed using a UPLC-ESI-MS/MS system (UPLC, SHIMADZU Nexera X2, https://www.shimadzu.com.cn/; MS, Applied Biosystems 4500 Q TRAP, https://www.thermofisher.cn/cn/zh/home/brands/applied-biosystems.html), and LIT and triple quadrupole (QQQ) scans were acquired using a triple quadrupole-linear ion trap mass spectrometer. Relative quantification of each flavonoid was conducted in multiple reaction monitoring (MRM) experiments. The analytical conditions were as follows, UPLC: column, Agilent SB-C18 (1.8 µm, 2.1 mm × 100 mm); the mobile phase consisted of solvent A (pure water with 0.1% formic acid) and solvent B (acetonitrile with 0.1% formic acid). In this study, 0.1 g AR samples in three biological replicates of each treatment were analyzed. The self-built Metware Database and metabolite information in the public database were used to determine the content of flavonoids, including 282 flavonoid-related metabolites: 10 chalcones, 29 flavanones, 20 anthocyanins, 26 flavanols, 13 flavanonols, 67 flavones, 77 flavanols, 11 proanthocyanidins, etc. (Supplementary file 1).

Anthocyanins metabolic class was detected using a UPLC-ESI-MS/MS system (UPLC, ExionLC™ AD, https://sciex.com.cn; MS, Applied Biosystems 6500 Triple Quadrupole, https://sciex.com.cn). Two anthocyanin metabolomic assays were conducted in this study. In the first assay, a total of 0.05 g with three sample replicates for each treatment was used. In UPLC-ESI-MS/MS assays of anthocyanin metabolites, linear ion trap (LIT) and triple quadrupole (QQQ) scans were acquired using a triple quadrupole-linear ion trap mass spectrometer. The analytical conditions were as follows, UPLC: column, Waters ACQUITY BEH C18 (1.7 µm, 2.1 mm × 100 mm); solvent system, water (0.1% formic acid): methanol (0.1% formic acid); gradient program, 95:5 V/V at 0 min, 50:50 V/V at 6 min, 5:95 V/V at 12 min, hold for 2 min, 95:5 V/V at 14 min; hold for 2min; flow rate, 0.35 mL/min; temperature, 40°C; injection volume, 2 μL. HPLC-grade methanol (MeOH) was purchased from Merck (Darmstadt, Germany). MilliQ water (Millipore, Bradford, USA) was used in all experiments. Formic acid was purchased from Sigma-Aldrich (St Louis, MO, USA). Anthocyanins were analyzed using scheduled multiple reaction monitoring (MRM). Data acquisitions were performed using Analyst 1.6.3 software and Multiquant 3.0.3 software (Sciex). In this study, anthocyanins were searched in the Metware database, which includes 44 anthocyanin-related metabolites: 35 anthocyanins, 6 flavonoids, and 3 proanthocyanidins (Supplementary file 2). The contents of 28 anthocyanin-related metabolites were determined by their standard substances, and the other 16 metabolites were semi-quantified using delphinidin-3,5-O-diglucoside. All of the standards were purchased from isoReag (Shanghai, China). The second metabolomic assays were conducted using three newly produced red poplar ARs. Same as the first assay, 0.05 g ARs samples were used in the second assay, however, the sample solution was 100 times diluted and then analyzed in UPLC-ESI-MS/MS system. A total of 10 anthocyanins, 2 flavonoids, and 4 proanthocyanidins were analyzed in the second assay. All these 16 metabolites were absolute quantified by their standard substances (Supplementary file 3).

A variety of statistical analysis methods, including principal component analysis (PCA), hierarchical cluster analysis (HCA), Pearson correlation coefficients (PCCs) and orthogonal partial least squares-discriminant analysis (OPLS-DA), were employed to distinguish the overall difference in metabolic profiles between groups and detect metabolite differences between groups. The relative importance of each metabolite to the PLS-DA model was checked using the variable importance in the projection (VIP) parameter. Significantly regulated metabolites between groups were determined by VIP ≥ 1 and fold change ≥ 2 or ≤ 0.5 or by VIP ≥ 1 for unique metabolites only in one group. Identified metabolites were annotated using the KEGG compound database (http://www.kegg.jp/kegg/compound/), and annotated metabolites were then mapped to the KEGG pathway database (http://www.kegg.jp/kegg/pathway.html). Pathways with significantly regulated metabolites were then subjected to MSEA (metabolite set enrichment analysis), and their significance was determined based on hypergeometric test P values.

### 2.4 Transcriptomic analyses

Corresponding to metabolomic assays, three Vso samples and three S-Vso samples were also analyzed in transcriptomic analysis. Total RNA of approximately 1.0 g ARs was extracted and used for RNA sequencing and RT-PCR validation. A total amount of 1 µg RNA per sample was used for cDNA Library Preparation and RNA Sequencing by Metware Biotechnology Co., Ltd. RNA sequencing was executed using NEBNext® UltraTM RNA Library Prep Kit for Illumina® (NEB, USA) following the manufacturer’s recommendations. The library preparations were sequenced using the Illumina HiSeq platform, and 125 bp/150 bp paired-end reads were generated. After removing low-quality (Q ≤ 20) and adapter reads, clean reads were assembled using reference genome sequence data with HISAT2. The *P. alba* var. *pyramidalis* genome sequence v1.0 (https://bigd.big.ac.cn/search/?dbId=gwh&q=GWHAAEP00000000) was used as the reference genome in this study. Gene expression was measured in fragments per kilobase of transcript per million fragments mapped (FPKM). DESeq2 was applied to identify differentially expressed genes (DEGs) between two groups, with FDR < 0.05 and fold change ≥ 2 or ≤ 0.5; for unique genes, FDR < 0.05 was used as the threshold for DEGs. Enrichment analysis was performed based on the hypergeometric distribution test. For KEGG, enrichment analysis was performed to determine metabolic pathways.

### 2.5 Combined analysis of the transcriptome and metabolome

The Pearson correlation algorithm was used to calculate the correlation between gene expression and metabolite response intensity data. DEGs and DEMs were selected to draw a correlation heatmap and correlation matrix. According to the results of association analysis between different genes and different metabolites, metabolome and transcriptome relationships were investigated to reveal the genes and metabolites playing crucial roles in pigment accumulation. Differentially expressed genes and metabolic pathways were also analyzed, and their common pathway information was mapped to KEGG pathways.

### 2.6 RT-qPCR validation

The expression of DEGs identified by the transcriptomic analysis and the expression of the key genes involved in flavonoid/anthocyanin biosynthesis was validated by RT qPCR using a FastKing RT Kit from Tiangen Co. (Beijing). The RT-qPCR primers were designed with National Center for Biotechnology Information Primer BLAST tools (available online: http://www.ncbi.nlm.nih.gov/tools/primer-blast/) and listed in Supplementary file 4. According to the previous reference gene selection method (Zhao et al. 2017), V-type proton ATPase catalytic subunit A (VHA-A, Potri.010G253500) was selected as the best reference gene for RT-qPCR validation from transcriptomic data for ARs. Relative transcript levels of target genes were calculated using the ^2-ΔΔCt^ formula (Livak and Schmittgen 2001). All RT-qPCR analyses were performed with four biological and 3 technical replications.

## 3. Results

### 3.1 Morphological characteristics of pathogen-induced adventitious roots

Phloem girdling-pathogen inoculation methods are novel methods that we developed for the physiological and pathological study of tree canker diseases (Xing et al. 2020). In a previous study, we reported allometric growth of phloem and xylem located on poplar stems above the girdling-inoculation site (Xing et al. 2020). In addition to this allometric growth, many ARs sprouted out from the new secondary phloem above the girdle or inoculation rings in *B. dothidea-* and *V. sordida*-inoculated poplar stem (Figure 1). As shown in Figure 1A-C, the new ARs (wrapped by PE film during AR formation) were always coral red or rosy in color. With variation based on experiment and season, 60-95% of pathogen-inoculated poplar stems developed ARs (unpublished data). The number of fibrous roots in the AR structure also changed in different experiments; however, one AR structure was able to develop 10 or even more fibrous roots, with the average diameter reaching 2-3 mm (data not provided). In contrast, no or only a few ARs (or a few fibrous roots) were observed in the girdling-cambium removal poplar stems (Figure 1D-E). Moreover, callus formation on the girdling rings was the typical characteristic (Figure 1F). Therefore, our observations suggest that poplar canker pathogens induced or promoted the formation of ARs on the poplar stems.

**Figure 1.**
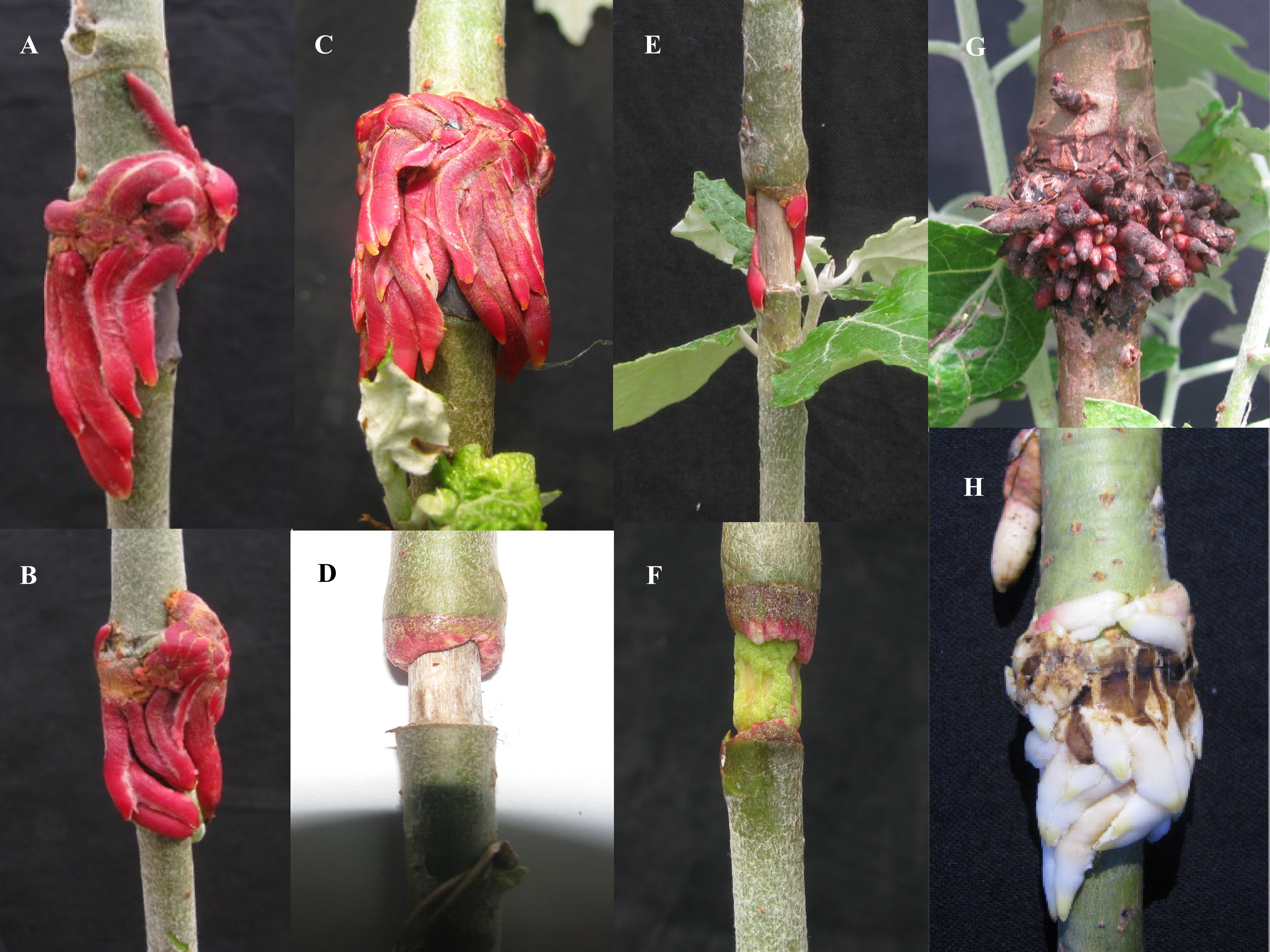
Canker pathogen inoculation induces the formation of adventitious roots (ARs) on poplar stems. AR formation under sunlight conditions (A-G): phloem-girdling, inoculated by pathogen *Valsa sordida* isolate C (A and B); phloem-girdling, inoculated by-pathogen *Botryosphaeria dothidea* isolate CZA (C); phloem-girdling and cambium removal, without pathogen inoculation, no or few AR formed (D and E); only phloem-girdling, callus tissue, but no or few ARs were observed (F); phloem girdling, *B. dothidea* inoculation, exposed to air for one week, with lignified and brown fibrous roots (G). AR formation under shaded conditions: phloem girdling, *B. dothidea* inoculation, and milky white ARs were observed (H). All the ARs or callus tissue were observed at 20-35 days after inoculation or girdling treatment.

The microscopic features of the ARs developed were as follows. Coral red or rosy pigments were observed in sunlight-exposed ARs (Figure 2A-C), and microscopy of AR sections and protoplasts showed the pigments to be water soluble and present in the vacuole of cells and not in the extracellular space (Figure 2F-G). Red cells were found in the epidermis of the outer surface of fibril roots and continuously or discontinuously distributed (Figure 2A-C). Red cells were also found in the cortex of fibril roots (approximately 35-45% of all cells), as clustered as some small vertical patches or discretely distributed; however, few or no red cells were found on the inner surface of the fibril root or the cortex near the light-exposed side. The color of the root tip of fibril roots in sunlight-exposed poplar ARs was always white or creamy yellow, and microscopy also showed that none or few red cells were present in the root tip and apical meristem of fibril roots in the newly developed ARs (approximately 15-20 days old). However, the whole fibril roots became red, and some red cells were observed in the root tip in mature ARs (25 days old) (Figure 1A-C; Figure 2A-C).

**Figure 2.**
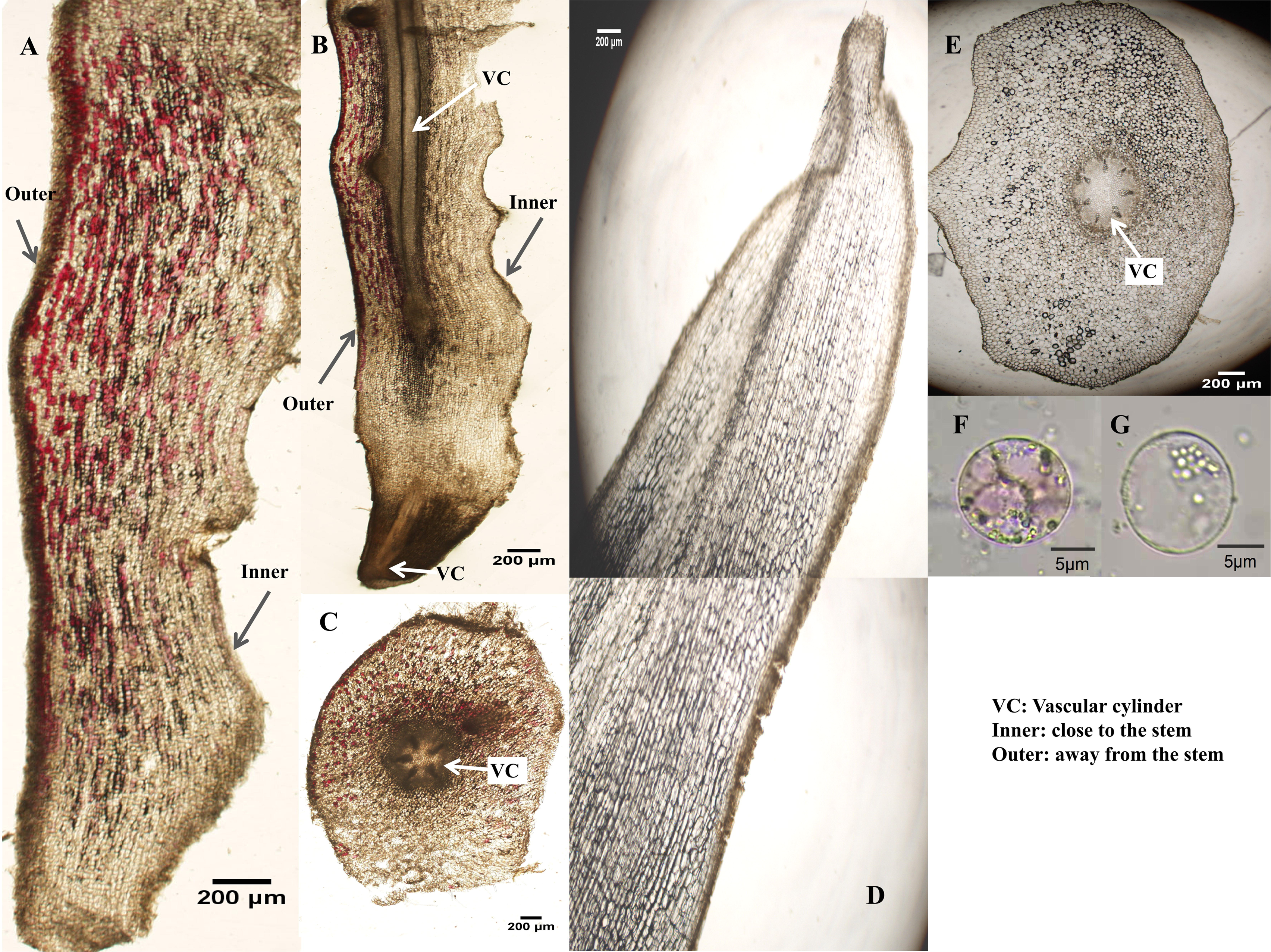
Distribution of pigment cells in ARs induced by the canker pathogen *Valsa sordida* on poplar stems. Microscopic observation of ARs formed under sunlight conditions (A-C) and under shaded conditions (D and E). Longitudinal section of root tips of ARs (A, B, and D) and transverse section root tips of ARs (C and E). Morphology of representative protoplasts derived from *Botryosphaeria dothidea*-induced ARs (F for colored protoplast and G for colorless protoplast). VC, vascular cylinder; Inner, the surface close to the poplar stem; Outer, the surface away from the poplar stem.

### 3.2 Influence of shading treatment on AR formation and pigment accumulation

The results illustrated that shading treatment did not influence the AR formation induced by pathogen inoculation on poplar stems (Figure 3A-B), the shape of ARs, or the number of all fibrous roots produced by each sapling. The length of every fibrous root was insignificant between sunlight and shading treatments (t-test, P >0.05, n ≥ 14) (Figure 3A; Supplementary file 5). However, pigment production was significantly inhibited in poplar ARs in shading conditions (Figure 1H). ARs formed under shaded conditions were always white in color, and the color of their root tip was always creamy yellow or canary yellow; moreover, consistent with the external appearance of ARs, no red cells were detected under the microscope in shaded poplar ARs (Figure 2D-E). Therefore, microscope observations also indicate that shading treatment inhibited pigment accumulation in poplar ARs. In other words, the results suggest that sunlight promotes pigment formation in ARs.

**Figure 3.**
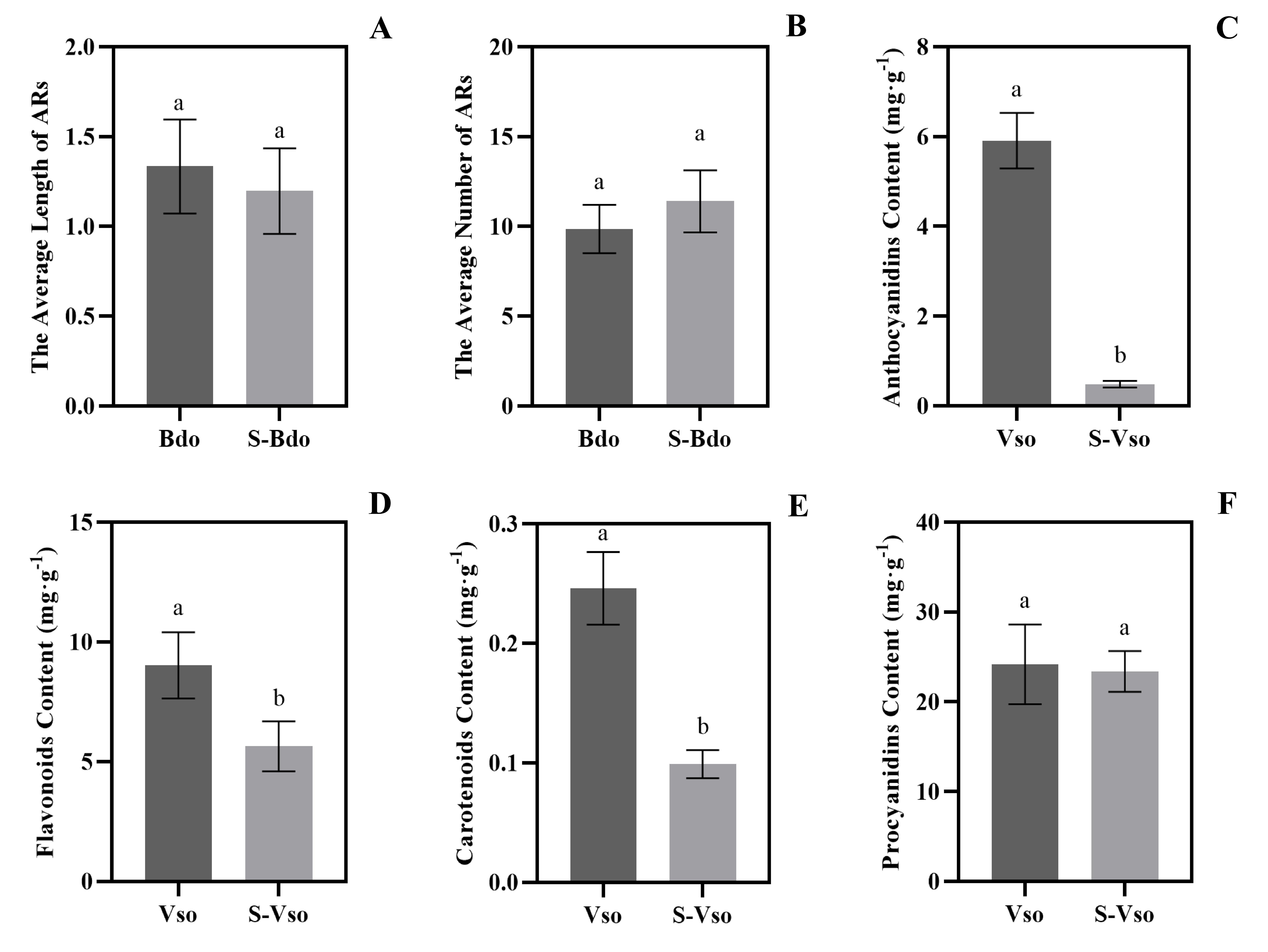
Length and number, total contents of anthocyanins, carotenoids, flavonoids, and proanthocyanidins of poplar ARs induced by canker pathogens. The average length (A) and number (B) of fibrous roots in every AR induced by *Botryosphaeria dothidea*. Contents of anthocyanins (A), flavonoids (C), carotenoids (B), and proanthocyanidins (D) in fibrous roots in every AR induced by *Valsa sordida*.

### 3.3 Pigment determination

As shown in Figure 3C-E, the content of anthocyanins in the Vso sample was 5.91 mg/g, 12.23 times that in S-Vso; the content of total flavonoids were 9.03 and 5,64 mg/g, in Vso and S-Vso, respectively; for carotenoids, the content was 0.25 and 0.99 mg/g (in Vso and S-Vso sample, respectively). The results showed that the total flavonoids, anthocyanins, and carotenoids significantly accumulated in Vso samples (t-test, p <0.05); however, carotenoids might not be the major colorant in poplar ARs for their relatively low content. In addition, though the content of proanthocyanidins (24.18 and 23.37 mg/g, in Vso and S-Vso, respectively) were higher than that of anthocyanins and flavonoids, there was no difference between Vso and S-Vso (Figure 3F, Supplementary file 6).

### 3.4 Metabolomic analysis

#### 3.4.1 Metabolomic analysis of flavonoid metabolites

In this study, OPLS-DA was used to identify differentially accumulated metabolites of flavonoid metabolic classes (DAFs) in poplar ARs. A total of 110 (93 up- and 17 downregulated) DAFs were identified (Supplementary file 1) in sunlight-exposed ARs (Vso sample), and the number of upregulated DAFs was significantly higher than that of downregulated DAFs (chi-square test, P < 0.01), suggesting sunlight significantly induce the accumulation of flavonoid metabolites. To simplify the description, all 110 DAFs were divided into three groups: High-, Middle- and Low-DAF, according to their relative intensity in sunlight-exposed ARs (Figure 4). The High-DAF group included 5 flavonols (rhamnetin-3-O-glucoside, F054; isorhamnetin-7-O-glucoside, F053; isorhamnetin-3-O-glucoside, F052; tamarixetin-3-O-(6’’-malonyl)-glucoside, F073; isorhamnetin-3-O-(6’’-malonyl)-glucoside, F072) and 2 flavones metabolites (nepetin-7-O-alloside, F055) and nepetin-7-O-glucoside, F056); the relative intensity of these metabolites are at least 2 times of that in Middle-DAF group. In addition to 7 flavonols and 7 flavones, the Middle-DAF group also included 4 anthocyanins (pelargonidin-3-O-glucoside, F021; cyanidin-3-O-glucoside (F034); delphinidin-3-O-galactoside, F046; and peonidin-3-O-glucoside, F047), and 4 another metabolite (1 tannin, chalcones and 2 flavonols). The relative intensity of the metabolites in the High- and Middle-DAF groups increased accumulated, while the accumulation of the first metabolites in the Low-DAF group, isosalipurposide (F025), was decreased in the sunlight-exposed poplar ARs (Figure 4). For the accumulation of Low-DAF metabolites was very low (only occupied at most ∼8.6% of the five top metabolites), these metabolites might play limited roles in the physiology of poplar ARs, therefore, this study suggested that sunlight induces the accumulation of flavonoids, mainly including subclass flavonols, flavones, and anthocyanins.

**Figure 4.**
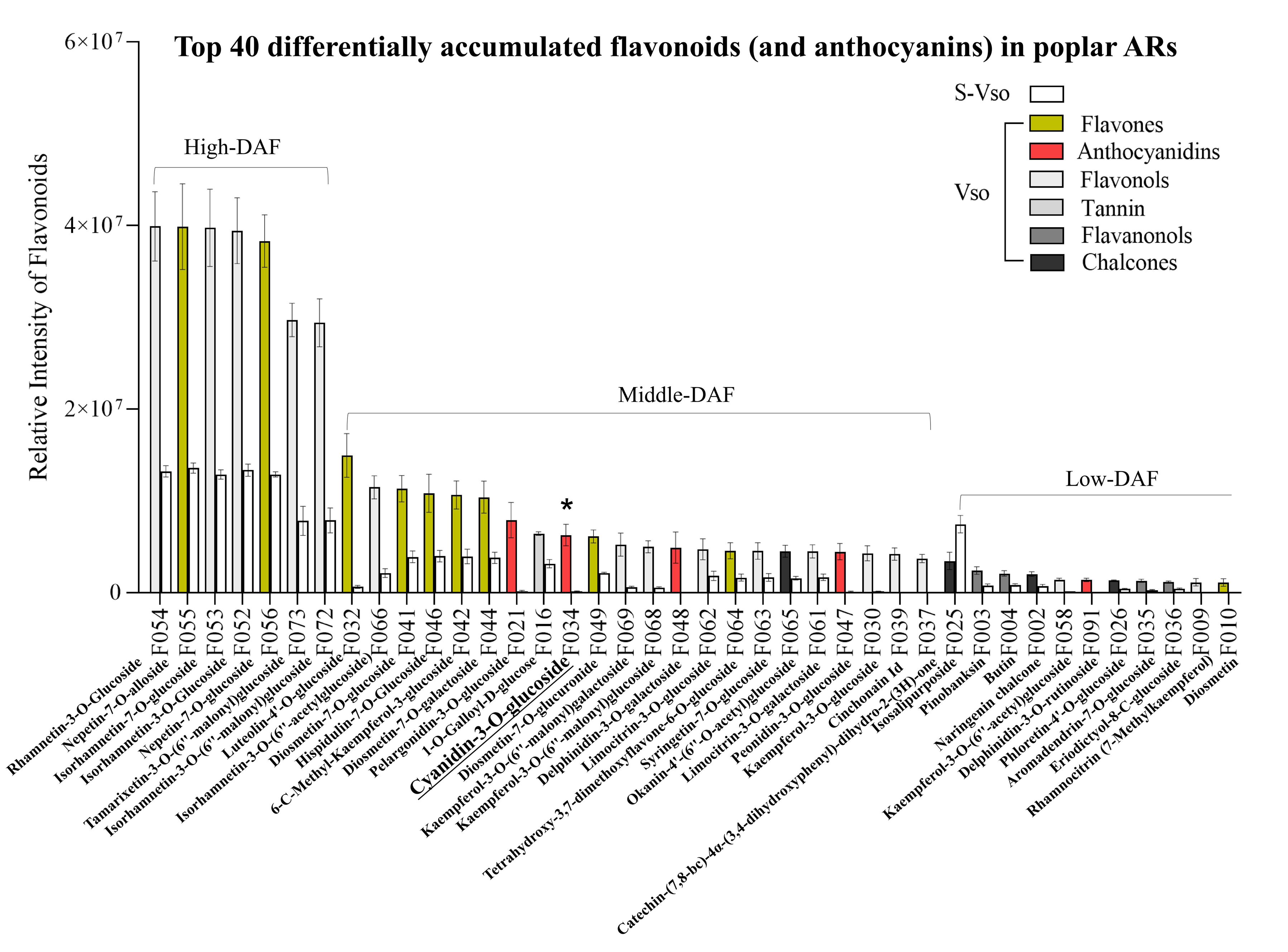
Top 40 differentially accumulated flavonoids and partial anthocyanins in sunlight-exposed poplar ARs. According to the relative intensity of flavonoid metabolites in the sunlight-exposed ARs, 110 differentially accumulated flavonoids (DAFs) were divided into three subgroups, high- (7 metabolites), middle- (22 metabolites), and low-DAF (81 but only the top 11 metabolites in this subgroup were exhibited here). The most abundant accumulated anthocyanin metabolite (cyanidin-3-O-glucoside) is highlighted with an asterisk. Different subgroups of flavonoid metabolites were exhibited in different colors; for example, red color represent the metabolites of anthocyanin group.

However, only flavone and anthocyanin metabolites are the color pigments of plants. To illustrate the coloration mechanisms of poplar ARs, categories and quantification of all differentially accumulated flavone and anthocyanin metabolites were further discussed. Nepetin-7-O-alloside (F055) and nepetin-7-O-glucoside (F056) (Figure 4, highlighted with brown color) exhibited the highest response intensity in HPLC-MS/MS assay, at 2.67-3.88 times that of the third flavone metabolites, luteolin-4’-O-glucoside (F032), and others. Pelargonidin-3-O-glucoside (F021) and cyanidin-3-O-glucoside (F034) are two subgroups of anthocyanins and the most highly expressed anthocyanins in the flavonoid assays (Figure 4, highlighted with red color); their response intensities were 19.8-15.7% of the value of the top 1 accumulated metabolite (F055). In addition, 2 flavones metabolites (diosmetin-7-O-glucuronide (F049) and tetrahydroxy-3,7-dimethoxyflavone-6-O-glucoside (F064)) and 2 anthocyanins (delphinidin-3-O-galactoside (F048) and peonidin-3-O-glucoside (F047)) showed response intensities similar to those of pelargonidin-3-O-glucoside and cyanidin-3-O-glucoside, but the relative intensity of the other flavones and anthocyanins metabolites were very low. Therefore, results suggest that the above metabolites all contribute to the color of poplar ARs; however, considering the color of poplar ARs in sunlight conditions, the 4 top accumulated anthocyanins metabolites might contribute more roles than flavones metabolites in the coloration of the red poplar ARs.

#### 3.4.2 Anthocyanin metabolomic analysis

A total of 44 metabolites (including 28 absolute quantified metabolites and 16 semi-quantified metabolites) were investigated in poplar ARs in anthocyanin assays in the first anthocyanin metabolomic assay (Supplementary file 2). However, the results of 28 absolute quantified metabolites (including 3 proanthocyanidins, 19 anthocyanins, and 6 flavonoids) were mainly reported in this study. Results showed that the quantification values of 26 metabolites were effectively quantified; however, the quantification values of the top 2 metabolites, procyanidin B3 (A107) and cyaniding-3-O-glucoside (A11), were both beyond their upper detection limit, and their calculated contents are 1313.50 ug/g and 717.36 ug/g (fresh weight), respectively. Results indicated that 21 metabolites were differentially accumulated in poplar ARs, including 17 anthocyanins (cyaniding-3-O-glucoside (A11), delphinidin-3-O-glucoside (A27), delphinidin-3-O-rutinoside (A29), etc.) and 2 flavonoids metabolites (naringenin (A39) and quercetin-3-O-glucoside (A41)) (Figure 5A). Besides procyanidin B3, procyanidin B1, and B2 were also highly accumulated in red poplar ARs, however, no difference was detected in three proanthocyanidins in the Vso and S-Vso comparisons (Supplementary file 2). After discarding colorless proanthocyanidin and flavonoid metabolites, 19 differential accumulated anthocyanins (DAAs) were ranked as their contents in the red poplar ARs (from the highest to the lowest). Besides the last three metabolites in the ranking, other 16 anthocyanins metabolites were increasingly accumulated in the red poplar ARs, suggesting that sunlight conditions increased the synthesis or accumulation of anthocyanins (Figure 5). The top 1, 2, and 4 accumulated anthocyanins were cyanidin-3-O-glucoside (A11), delphinidin-3-O-glucoside (A27), and pelargonidin-3-O-glucoside (A68), which was the first production of cyaniding, delphinidin, and pelargonidin metabolites synthesis.

**Figure 5.**
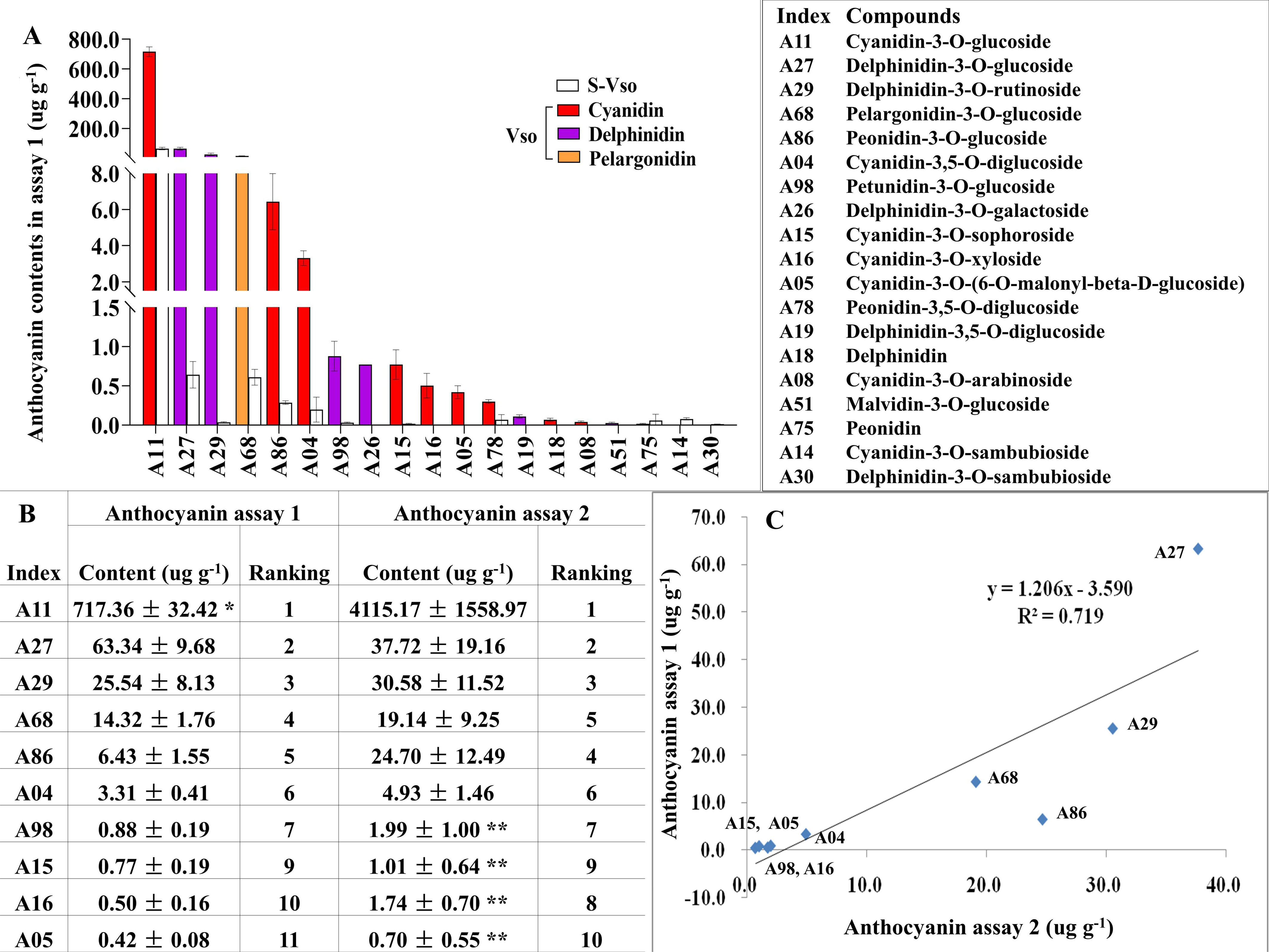
Differentially accumulated anthocyanins (DAAs) in sunlight-exposed poplar ARs. A total of 19 anthocyanin metabolites differentially accumulated in sunlight-exposed poplar ARs in the third anthocyanin metabolomic assay (A), the contents of all metabolites were determined by their authentic standards. The comparison of the contents of the top 10 accumulated anthocyanins detected in anthocyanin metabolomic assay 1 and 2 (B). Linear correlation analysis of the anthocyanin contents between anthocyanin assay 1 and 2 (C). The content value marked with an asterisk (*) represent the quantification of the anthocyanin beyond its upper limit of quantification, while the content value marked with double asterisk (**) represent the quantification of the anthocyanin beyond its lower limit of quantification. Cyanidin metabolites are highlighted with red color, delphinidin metabolites are highlighted with purple color, and pelargonidin metabolites are highlighted with brown color.

To determine the authentic contents of cyanindin-3-O-glucoside (A11) and procyanidin B3 in poplar ARs, the second anthocyanin metabolomic assay was conducted in three newly produced poplar ARs in sunlight conditions. The quantifications of 16 metabolites (including 10 anthocyanins, 4 proanthocyanidins, and 2 flavonoids) were determined using their standard substances (Supplementary file 3). Results showed that the content ranking of 10 anthocyanin metabolites detected in the second assay is almost identical to that in the first assay (Figure 5B). Moreover, cyanindin-3-O-glucoside (A11) is still identified as the highest accumulated anthocyanins in poplar ARs, which content is 4,115.17 ug/g (fresh weight), 5.73 times the first measured value (717.36 ug/g fresh weight) and 109.15 times of the top 2 metabolite delphinidin-3-O-glucoside (A27, 37.72 ug/g fresh weight). Besides the huge difference of the content in cyaniding-3-O-glucoside (A11) in two assays, the contents of other anthocyanins were at the same level, for example, delphinidin-3-O-rutinoside (A29), pelargonidin-3-O-glucoside (A68), Cyanidin-3,5-O-diglucoside (A04), etc. A linear correlation relationship of the contents of other 9 anthocyanins (cyandin-3-O-glucoside (A11) was discarded) was detected between the first and second assays (y=1.206x-3.590, R^2^=0.719, Figure 5C). However, only the top 6 anthocyanin metabolites were accurately quantified in this assay because the MS quantification values of last four anthocyanins were beyond their lower limit of quantification, which might be caused by the over-dilution (100 times) of injection solution in MS assay. Additionally, results also showed that procyandin B1 and B3 are the highest accumulated proanthocyanidins, their contents are 6,326.34 ug/g and 4,245.11 ug/g, respectively.

Cyanidin-3-O-glucoside (A11) was also detected in metabolic assays of flavonoid class and noted as F034 with a relatively high response intensity (top 9 metabolites in all “real color-related” flavonoids and top 2 in anthocyanins) in poplar sunlight-exposed ARs. Thus, the flavonoid metabolome and two anthocyanin metabolomes suggested that cyanidin-3-O-glucoside is the most abundant anthocyanin and the most dominant colorant in poplar ARs. Moreover, results also showed that proanthocyanidin metabolites highly but insignificantly accumulated in poplar ARs.

#### 3.4.3 KEGG enrichment analysis of flavonoid and anthocyanin metabolomes

In this study, differentially accumulated flavonoid metabolic class (DAFs) and anthocyanins (DAAs) were enriched in KEGG pathways (Supplementary file 7). Twelve DAFs were enriched in the “anthocyanin biosynthesis” pathway; in addition to F085 and F087, 10 of 12 DAFs were upregulated in Vso saplings (Supplementary file 7). In anthocyanin analysis, 20 DAAs were enriched in the “anthocyanin biosynthesis” pathway, and 18 of these DAAs were upregulated in Vso. All 12 DAFs are also related to anthocyanin metabolism; in addition to metabolites pelargonidin-3,5-O-diglucoside (F085/A59) and cyanidin-3-O-rutinoside (F087/A12), the majority of 12 DAFs showed a similar regulatory pattern in the metabolic analyses of flavonoid and anthocyanin metabolites.

In plants, all anthocyanin derivatives derive from 3 pathways: the cyanidin, delphinidin, and pelargonidin pathways. In our study, the accumulation of almost all metabolites increased in poplar ARs formed in sunlight conditions. For example, the accumulation of 8 of 9 anthocyanins in the cyanidin pathway and 5 of 6 derivatives in the delphinidin pathway were induced, particularly, the amount of cyanidin-3-O-glucoside (A11) was greatly increased in the poplar sunlight-exposed ARs. The results suggest that light treatment increases the accumulation of almost all anthocyanin metabolites in the cyanidin-, delphinidin- and pelargonidin-based pathways. However, considering the content of anthocyanins, light exposure greatly increases the content of cyanidin-base anthocyanins in poplar ARs.

### 3.5 **Transcriptomic analysis and RT-qPCR validation**

#### 3.5.1 Transcriptome analysis reveals the differently expressed genes involved in the flavonoid biosynthesis pathways

Compared with the poplar shaded samples (S-Vso) a total of 803 differentially expressed genes (DEGs) were detected in the sunlight-exposed ARs (Vso) (Supplementary file 8). These DEGs were enriched in 6 light- or photosynthesis-related pathways based on KEGG enrichment analysis, such as photosynthesis-antenna proteins, photosynthesis, circadian rhythm-plant, porphyrin and chlorophyll metabolism, and carbon fixation in photosynthetic organisms (P < 0.05; Supplementary file 7). The results showed the majority of DEGs to be upregulated in the sunlight-exposed ARs at the genome-wide level (588 upregulated and 215 downregulated; chi-square test, P < 0.01). In particular, all 21 genes in the photosynthesis-antenna protein pathway (encoding 7 subgroups of the LHC complex), all 40 genes in photosynthesis (encoding proteins or subgroups of PI, PII, electron transporter, and F-type ATPase), and 19 of 21 DEGs in circadian rhythm were upregulated in the sunlight exposed ARs (Supplementary file 8). The results highlight that expressions of light- or photosynthesis-related genes are promoted in sunlight-exposed poplar ARs.

In this study, the expression of 13 flavonoid-related DEGs was detected in poplar ARs, including ten upregulated genes, such as genes encoding C3H (C3-hydroxylase), HCT (hydroxycinnamoyl-CoA shikimate/quinate hydroxycinnamoyl transferase), FLS (flavonol synthase/flavanone 3-hydroxylase), LAR (leucoanthocyanidin reductase), and BZ1 (anthocyanin 3-O-glucosyltransferase), respectively. The results showed that 4 BZ1 genes (such as the PAYG030937 gene) upregulated expressed in poplar ARs under sunlight conditions; however, two transcripts of HCT and 1 ANS (anthocyanin synthase) downregulated expressed. Additionally, 1 IF7MAT gene (isoflavone-7-O-glucoside-6’’-O-malonyltransferase) and 3 FG2 genes (flavonol-3-O-glucoside L-rgamosyltransferase) increasingly expressed in flavones and flavonoid biosynthesis pathway (ko00944); 1 IF7MAT, 1 HIDH (2-hydroxyisoflavanone dehydratase), and 1 PTS gene (pterocarpan synthase) increasingly expressed in isoflavonoid biosynthesis pathway (ko00943), these results suggested the high accumulation of flavonols and flavones metabolites in the sunlight-exposed poplar ARs.

In transcriptomic analysis, 69 differentially expressed TFs were identified between Vso and S-Vso (Supplementary file 8). Among these TFs, 3 MYB113 (R2R3_MYB) genes (PAYG035595-7, homology with *AtMYB113*), 3 bHLH genes (PAYG007639 and PAYG031606, homology with *AtbHLH38*; PAYG017891, homology with *AtbHLH16*), and 3 WD40 domain protein genes (PAYG025818, PAYG012381 and PAYG018605), 1 COP1 gene (encoding Ring-finger protein CONSTITUTIVE PHOTOMORPHOGENIC1, PAYG018730), 1 PIF3 gene (phytochrome-interacting factor 3, PAYG017891), and 1 HY5 gene (ELONGATED HYPOCOTYL5, PAYG035590) were upregulated in sunlight exposed poplar ARs. While 3 bHLH genes (PAYG002116 and PAYG25886, homology with *At*bHLH111; PAYG033885, homology with *At*bHLH38) and 1 SPL gene (encoding squamosa-promoter binding protein-like protein) were downregulated.

#### 3.5.2 RT-qPCR validation of transcriptomic analysis

In this study, 30 DEGs identified by the transcriptomic analysis were validated by RT-qPCR using a FastKing RT Kit from Tiangen Co. (Beijing). Expression of a total of 30 DEGs, including 12 genes encoding ANS, BZ1, FLS, LAR, HCT, FG2, IF7MAT, etc., and TFs related to the biosynthesis of flavonoids, encoding MYB113 and SPL (squamosa promoter-binding-like protein), were validated by RT-qPCR. As illustrated in Supplementary file 9, in addition to the FG2 gene (PAYG014565, data not shown), the other 29 DEGs were validated by RT-qPCR assays. Specifically, genes encoding BZ1, FLS, LAR, HCT, FG2, and IF7MAT were upregulation expressed while genes encoding ANS and SPL were downregulation expressed in the sunlight-exposed poplar ARs. Moreover, the expression of 29 of 30 DEGs based on RT-qPCR positively correlated with that based on transcriptome analysis (y=1.58 x-0.003397, R^2^ =0.72, P < 0.01) (the last chart in Supplementary file 9).

### 3.6 Light-induced color formation in shaded poplar ARs

To validate the above conclusion, white ARs produced under shaded conditions were exposed to sunlight for three days. As shown in Figure 6A, at 4 hours after sunlight exposure, red pigments gradually accumulated on the surface of shaded poplar ARs. One day after light exposure, the color of the AR fibrils became coral red or rosy, similar to the color of the ARs induced under sunlight conditions for two weeks or even a long time (Figure 1). Whether sunny or cloudy weather, red pigments can produce on the surface of ARs after removing the aluminum foil. Moreover, we observed that the change in poplar AR color was faster on sunny days than on cloudy days.

**Figure 6.**
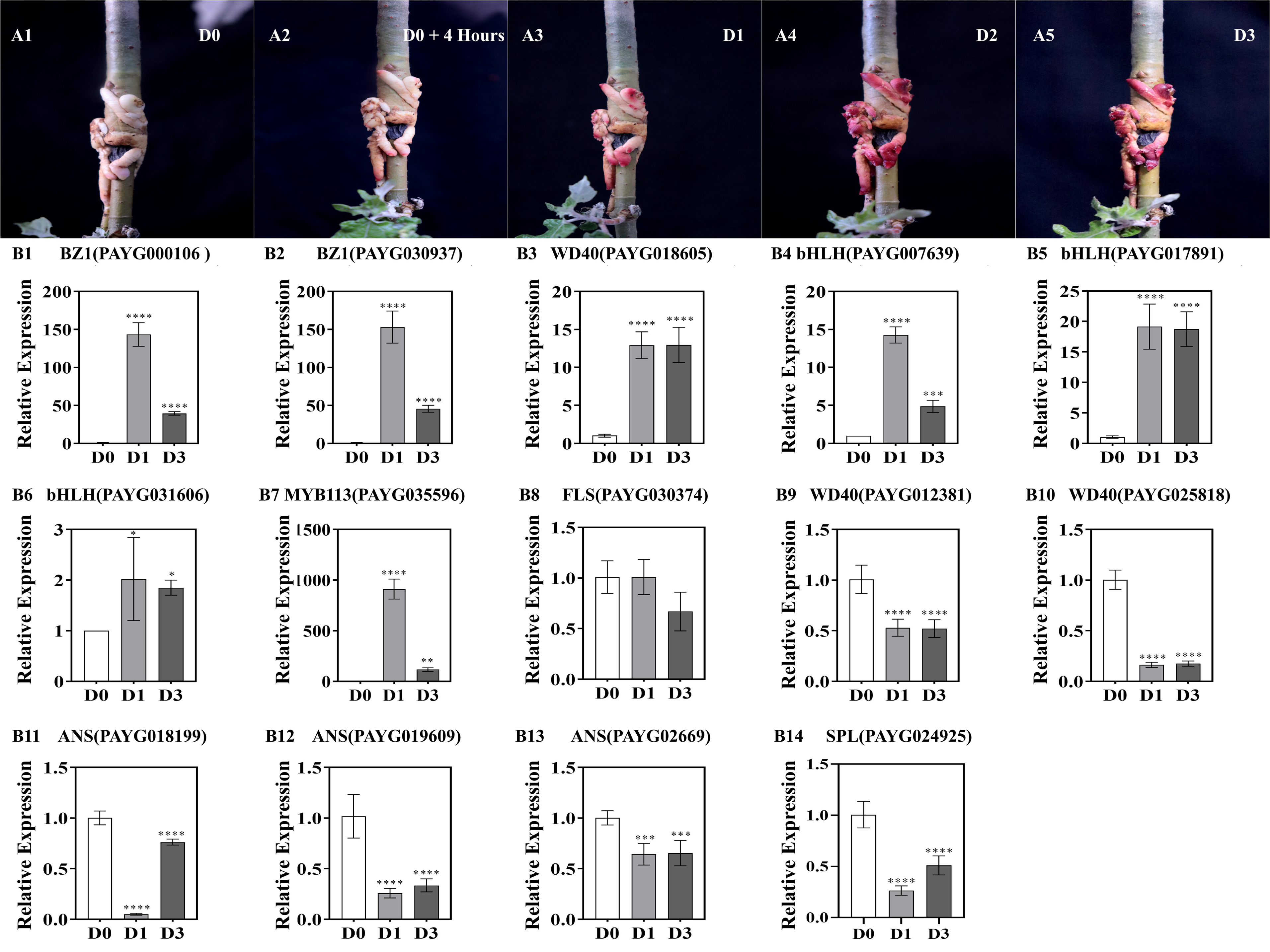
Light induces pigment formation and expression of anthocyanin biosynthesis-related genes in shaded poplar ARs. Pigment accumulation in shaded poplar ARs at three days after exposure to sunlight (A). Expression of 14 genes related to anthocyanin biosynthesis (B). B1-B2: BZ1 (Bronze 1, anthocyanin 3-O-glucosyltransferase); B3, B9-B10 (WD40); B4-B6 (bHLH); B7 (MYB113); B8 (FLS, flavonol synthase/flavanone 3-hydroxylase); B11-B13 (ANS, anthocyanin synthase); B14 (SPL, squamosa-promoter binding protein-like proteins). The significantly expressed genes were noted with an asterisk or asterisks (t-test, *, p < 0.05; **, p <0.01).

We detected the expression of genes involved in anthocyanin biosynthesis, such as 6 structural genes (1 FLS, 3 ANS, and 2 BZ1 encoding genes), and 8 transcript factor genes (1 MYB113, 2 bHLH38 and 1bHLH16, 3 WD40, and 1 SPL encoding gene) in shaded poplar ARs exposed to sunlight. As shown in Figure 6B1-B7, the expressions of 2 BZ1, 1 WD40, 3 bHLH, 1 MYB113, and 1 WD40 (PAYG018065) encoding genes were upregulated at 1 and 3 days after sunlight exposure, consistent with the regulatory patterns of these genes in transcriptomic analysis. Moreover, results showed that the relative expression of 2 BZ1 genes (PAYG030937 and PAYG000106) increased about 150 times at 1 day-after-exposure (Figure 6B1-B2), which might be the reason for the rapid biosynthesis of anthocyanins after light exposure. On the other hand, the expression of 3 ANS and 1 SPL gene was inhibited at 1 and 3 days after sunlight exposure (Figure 6B11-B14), also consistent with the regulatory patterns of these genes in transcriptomic analysis. Aside from 1 FLS gene (PAYG030374, not changed after sunlight exposure) and two WD40 genes (PAYG012381 and PAYG025818; both inhibited at 1, 3 days after sunlight exposure), this result is well consistent with our transcriptomic results (11 in 14 validated genes): the expression of genes encoding BZ1 is co-upregulated with the expression of genes combining “MBW” complex, while the expression of genes encoding ANS and SPL were downregulated at sunlight-exposure conditions in poplar ARs.

## 4. Discussion

Adventitious roots are plant roots that form from any non-root tissue and can be produced during normal development or in response to stress conditions like flooding, nutrient deprivation, and wounding (Steffens and Rasmussen 2016). Some woody perennials, such as willow and poplar species are more likely to form ARs due to the preformed AR initials (albeit dormant) already existing in their stems (Bellini et al. 2014). Under tissue culture conditions, in soil, or in cultivation substrates, phytophormones (mainly auxin) stimulate ARs developed from the end of cuttings and leaves. This is one reason for the largest cultivation area of poplar plantations developed in China in the world. Moreover, symbiotic interactions between plants and beneficial microorganisms, mainly mycorrhizal fungi and rhizobial bacteria, can affect the root system architecture (general root growth, length of primary and lateral roots (LR), LR number, and positioning) and stimulate ARs development in hypocotyl or stem cuttings (Bellini et al. 2014). For instance, arbuscular mycorrhizal fungi, such as *Glomus intraradices*, or their secretions or cultures, can stimulate LR development, induce root initiation–related genes and establish a root development program in *Medicago truncatula* (Maillet et al. 2011; Oláh et al. 2005). Ectomycorrhizal fungi like *Laccaria bicolor* and *Tuber melanosporum* can trigger LR formation before colonization in poplar (*Populus tremula* × *P. alba*) and *Cistus incanus*, respectively, as well as in *Arabidopsis* (Felten et al. 2009; Splivallo et al. 2009). Additionally, the bacterial infection of the pathogen *Agrobacterium rhizogenes* can induce “hairy roots” from any vegetative part of the plants by transferring a bacterial *rolB* (*root oncogenic locus* B) gene into the plant host cells (Coudert et al. 2013), providing an efficient transgenic method in plant biotechnology. However, no research about fungal pathogens that induce or promote root development or AR formation was reported.

In this study, colorful and plenty of poplar ARs were observed in our experiments (Figure 1, Figure 2, and Figure 6), using the novel girdling-inoculation method (Li et al. 2019; Li et al. 2021; Xing et al. 2020; Xing et al. 2022) which only used in our physiological and pathological researches of poplar stem canker diseases. To our best knowledge, the formation of poplar ARs induced by fungal pathogens was a novel allometric phenomenon in woody species. The mechanisms of pathogens induce the formation, the epigenetic response of ARs and the derived explants to pathogens, and the potential use in agriculture and forestry were investigated in our laboratory. Here, mechanisms of flavonoid/anthocyanin biosynthesis, and then the coloration mechanism in pathogen-induced poplar ARs under sunlight exposure are reported.

### 4.1 Sunlight exposure changes the accumulation and metabolic flux of flavonoids in poplar ARs

In this study, chemical and metabolomic assays illustrated that sunlight exposure increases the accumulation of metabolites in class flavonols, flavones, and anthocyanins. Flavonols and flavones are the two major accumulated classes of flavonoids in the sunlight-exposed poplar ARs, which occupied 67 metabolites in all 110 differentially accumulated flavonoids (DAFs) and all 13 top accumulated DAFs (including all 7 members of the High-DAF group and top 6 members of Middle-DAF group (Figure 4; Supplementary file 1). Anthocyanin is the third high accumulated flavonoid-related metabolite in poplar ARs: four kinds of highly accumulated anthocyanins were detected and validated in both flavonoids and anthocyanins metabolome, while metabolome showed that 29 in 34 anthocyanins metabolites increasing accumulated in the sunlight-exposed poplar ARs (Figure 4, Supplementary file 1, 2).

In plants, the flavonoid/anthocyanin biosynthesis pathway produces many flavonoids metabolites (such as chalcone, flavanones, flavanonols, dihydroflavonols, leucoanthocyanidins, anthocyanidins, and anthocyanins); however, the other flavonoids classes (flavones, flavonols, isoflavones, and proanthocyanidins) produced from different branching pathways (Nabavi et al. 2020). Anthocyanin synthase (ANS) and BZ1 (Bronze 1) are the last two enzymes involved in anthocyanins biosynthesis, which catalyze the conversion of leucoanthocyanidin to anthocyanidins and then the glycosylation of anthocyanidins (Feng et al. 2021; Wilmouth et al. 2002). Coexpression of gene ANS and BZ1 were observed in plants and positively associated with the biosynthesis of flavonoid/anthocyanin (Zhou et al. 2022; Liu et al. 2022b); moreover, the other genes involved in flavonoids biosynthesis (such as genes encoding PAL, C4H, CHS, F3H, ANR, F3’Hs, DFRs, LAR, GT1, etc.) were also significantly upregulated (Liu et al. 2022b), suggesting a de novo biosynthesis of flavonoids and anthocyanins in duckweed variety. In our recent study, aside from gene FLS, gene PAL, CHS, CHI, F3H, DFR, ANS, LAR, and ANR are all upregulated, suggesting a de novo biosynthesis of flavonoids in the BCMV (bean common mosaic virus) infected poplar leaves (Wang et al. 2023). However, one recent research illustrated that ANS genes negatively but BZ1 genes positively regulate the biosynthesis of anthocyanins in poplar 84K leaves (Yan et al. 2022). In this study, downregulation of the ANS gene was also observed in the sunlight-exposed poplar ARs, suggesting a block of metabolic flux at the leucoanthocyanidins position; therefore, no or fewer anthocyanidins can be produced from leucoanthocyanidins. Then, high accumulation of three anthocyanins (cyaniding-3-O-glucoside, A11; pelargonidin-3-O-glucoside, A68; and delphinidin-3-O-glucoside, A27) (Figure 7) were converted from the existing anthocyanidin compounds, catalyzed by the upregulation of BZ1 gene (the second chart in Supplementary file 9). In a word, the results in this study illustrated a direct, substrate-depended biosynthesis of anthocyanins. As the only differentially expressed gene in the flavonoid/anthocyanin biosynthesis pathway, obviously, the upregulated BZ1 gene plays a crucial role in the synthesis of anthocyanins, and in the coloration of poplar ARs. However, the mechanism of light exposure inhibited the expression of ANS (PAYG019609) in poplar plants needs further investigation.

**Figure 7.**
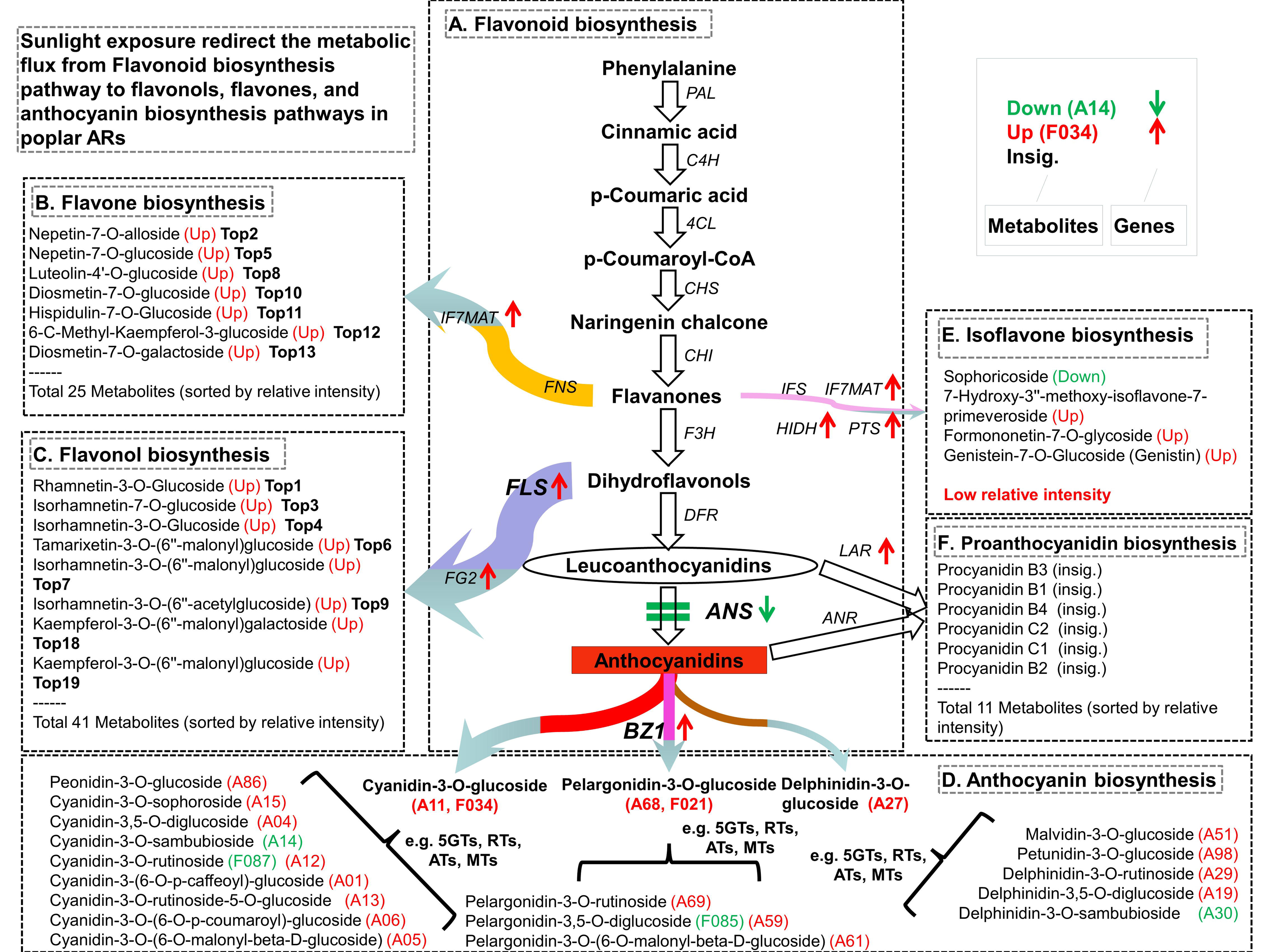
Sunlight exposure redirects the metabolic flux from the flavonoid biosynthesis pathway to flavonols, flavones, and anthocyanin biosynthesis pathways in poplar ARs. Integrated analysis of metabolomic and RNA sequencing illustrated that sunlight exposure changed the gene expression and redirected the metabolic flux of the flavonoid biosynthesis pathway in poplar ARs. The widest, indigo arrow represents the shift of metabolic flux from the flavonoid biosynthesis pathway to the flavonols biosynthesis pathway, catalyzed by the increasing expressed FLS gene; the second widest, orange arrow represents the shift of metabolic flux from the flavonoid biosynthesis pathway to the flavone biosynthesis pathway. The double horizontal lines (in green) represent a block of metabolic flux from leucoanthocyanidins to anthocyanidins, which is caused by the down expression of ANS genes; meanwhile, the multiple arrows represent the direct biosynthesis of anthocyanins based on substrate anthocyanidins, catalyzed by the increased expression of BZ1 genes in sunlight conditions in poplar ARs. The red, up solid arrows represent the neighbor gene upregulation expressed; while the green, down solid arrows represent the neighbor gene downregulatory expressed. The width of the arrows represents the relative intensity of metabolic flux; for example, the widest, purple arrow represents the most abundant metabolic flux redirected to the flavonols pathway. A, the diagram of the flavonoid biosynthesis pathway; B, the branch pathway of the flavones biosynthesis and the most accumulated metabolites; C, the branch pathway of the flavonols biosynthesis and the most accumulated metabolites; D, the anthocyanin biosynthesis pathway and the detected metabolites in the anthocyanin metabolomic analysis; E, the branch pathway of the isoflavones biosynthesis and the differentially accumulated metabolites; and F, the branch pathway of proanthocyanidin biosynthesis and the top accumulated metabolites, insignificant accumulated between the shaded and sunlight-exposed poplar ARs. Three committed enzymes of biosynthesis of flavonols or anthocyanins, FLS (flavonol synthase/flavanone 3-hydroxylase), ANS (anthocyanin synthase), and BZ1 (Bronze 1, anthocyanin 3-O-glucosyltransferase), were in bold exhibited.

Flavonol synthase (FLS) is the key committed enzyme in the biosynthesis of class flavonols, which convert dihydroflavonols to the form of quercetin, kaempferol, and myricetin (Kou et al. 2023). Considering the high accumulation (the relative intensity, number of top accumulated metabolites, and total accumulated metabolites) of flavonols, the upregulation of gene FLS (Supplementary file 1, Supplementary file 9) suggested a redirection of metabolic flux into the flavonols branching pathway. Compared to the shade conditions, sunlight exposure promoted the accumulation of 37 flavonols metabolites, including 5 top accumulated metabolites in the High-DAF group (Figure 4). Though the expression of FNS, the crucial enzyme controlling metabolic flux to the flavones branching pathway, did not significantly change, metabolomic results illustrated that metabolomic flux redirected to the flavones branching pathway from the flavonoid/anthocyanin biosynthesis pathway (Figure 7; Supplementary file 1, 2). The accumulation of 25 flavones metabolites, including two flavones in the High-DAF group, significantly increased in sunlight-exposed poplar ARs (Figure 4). Moreover, chemical and metabolomic assays showed that proanthocyanidin is the most abundant accumulated flavonoids class; however, there was no difference in their contents or relative intensities between the sunlight-exposed and shaded poplar ARs (Figure 3F, Supplementary file 2). Therefore, this result illustrated sunlight exposure cannot influence the accumulation of proanthocyanidins, though the expression of gene LAR increased in sunlight-exposed poplar ARs.

### 4.2 Accumulation and coloration of anthocyanins in poplar ARs

Anthocyanins are the major colorants of flowers, fruits, buds, and young shoots, as well as of purple and purple-red leaves in autumn (Falcone Ferreyra et al. 2012; Kayesh et al. 2013; Mekapogu et al. 2020; Vaknin et al. 2005). Anthocyanins also accumulate and impart a red color to the roots or ARs of herbaceous plants, such as purple carrots, cassava roots, and *Raphanus sativus* (Chialva et al. 2021; Fu et al. 2022; Xiao et al. 2021), and accumulate in the ARs of the woody plant *Metrosideros excelsa* (Solangaarachchi and Gould 2001). In this study, chemical assays illustrated that flavonoids, anthocyanins, and carotenoids significantly accumulated in poplar ARs under sunlight exposure (Figure 3C-E), and then, the metabolomic analysis identified the accumulated flavonoids metabolites as flavonols, flavones, and anthocyanins (Figure 4).

However, flavonols are not the major colorant of poplar red ARs because of their spectral features (colorless), though they highly accumulated in poplar ARs. The flavones and carotenoid metabolites also are not the major colorant of poplar ARs for their yellow spectral features. Then, the coral red or rosy red color of poplar ARs (Figure 1A-C; Figure 6A), pigments distribution (Figure 2A-C), and the color stability in different pH conditions (Supplementary file 10) suggested that the red colors are related to anthocyanins metabolites. Finally, this speculation was validated in our flavonoids and anthocyanins metabolomic analysis (Figure 3C-D, Figure 4, and Figure 5). Anthocyanins can be divided into three subclasses, cyanidin-, pelargonidin-, and delphinidin-base metabolites. However, cyanidin-base anthocyanins were reported as the major colorant of plant fruits and seeds (Cevallos-Casals and Cisneros-Zevallos 2003; Khoo et al. 2017; Seeram et al. 2001). Especially, cyanidin 3-O-glucoside was reported as the major or one of the major anthocyanins metabolite in the different organs of woody plants, such as red-leaf of acer (Zhang et al. 2021), red callus of *Vitis davidii* (Lai et al. 2022), red anthers or leaves of poplar (Cho et al. 2016; Kim et al. 2021), and leaves of *AmRosea*1 transgenic 84K poplar (Yan et al. 2022). This study also illustrated that cyanidin-3-O-glucoside is the most abundant and absolutely dominant colorant in the sunlight-exposed poplar ARs, its content can up to 4115.17 ug/g in poplar red ARs. Moreover, flavonoid metabolomic assays showed that seven colorful flavones metabolites (especially for nepetin-7-O-alloside and nepetin-7-O-glucoside) highly accumulated in poplar ARs (Figure 4), suggesting flavones metabolites also contribute to the coloration of poplar ARs. In other words, both anthocyanins and flavones metabolites contribute to the red color of poplar ARs; however, anthocyanins play a predominant role in the coloration of sunlight-exposed poplar ARs.

### 4.3 Light regulates the accumulation of flavonoids/anthocyanins in poplar ARs

It is well known that light regulates biosynthesis of primary and secondary metabolites, such as flavonoids and anthocyanins (Zoratti et al. 2014a; Zoratti et al. 2014b), and the light quality (or spectra) as well as light intensity influence flavonoid and anthocyanin biosynthesis (Cominelli et al. 2008; Deng et al. 2012; Pan and Guo 2016; Wang et al. 2012; Xu et al. 2014). In this study, sunlight-exposure experiments were conducted to explore the influence of light on coloration and pigment biosynthesis in poplar ARs. The results showed sunlight induces a high accumulation of flavonols, flavones, and anthocyanins in the poplar ARs, resulting in a change of coloration and pigment contents (Figures 1, Figure 3, Figure 6).

Research shows that ARs have high sensitivity to sunlight exposure, and anthocyanin metabolites were rapidly produced after sunlight exposure (Solangaarachchi and Gould 2001). Our research also illustrated high photosensitivity in poplar ARs: some visible color alterations were observed in the white, shaded poplar ARs after 1-3 hours when they are exposed to sunlight (Figure 6A) In our epidermis girdling-inoculation experiments, many fibril roots became brown or even black and lignified when they were exposed to air (Figure 1G), breaking through the Parafilm^®^ package. Compared to this normal root morphology, the huddled morphology of poplar Ars (Figure 1A-F, H) might be due to the PE plastic wrapping. The rapid transcriptional activation of the flavonoid biosynthesis pathway is one trait of plants acclimating to high light, which results in the rapid accumulation of photoprotective and antioxidative flavonoids, such as flavonols and anthocyanins, in the leaf tissue (Araguirang and Richter 2022). Then, the upregulation of gene BZ1 and FLS suggested transcriptional activations of the flavonols and anthocyanin-related genes in poplar ARs. Moreover, compared to the de novo biosynthesis from phenylalanine or other upstream intermediates, the direct and anthocyanidins-dependent anthocyanin biosynthesis (which due to the downregulation of ANS and upregulation of BZ1), can produce photoprotective anthocyanins, provide light attenuation and antioxidation response to light stress after a short time (Zheng et al. 2019), and then increase the poplar acclimation to cope with sudden sunlight exposure stress.

### 4.4 Gene regulation in the flavonoid/anthocyanin biosynthesis in poplar ARs

Genes involved in flavonoid/anthocyanin biosynthesis are divided into two classes according to their role in metabolite biosynthesis: structural and regulatory genes (Jaakola et al. 2002; Koes et al. 2005). Structural genes encode the enzymes that produce anthocyanins from phenylalanine to cyaniding-, delphinidin-, and pelargonidin-base metabolites. Regulatory genes, for example, the R2R3-MYB (MYB113, MYB7) transcript factors, play key regulatory roles in flavonoid/anthocyanin biosynthesis and the main coloration determinant in plants (Cavagnaro et al. 2014; Chialva et al. 2021; Feng et al. 2021; Xu et al. 2019). However, in most cases, three transcript factors (R2R3-MYB, basic helix-loop-helix (bHLH), and WD40 repeat protein families) function together (consistently up- or down-regulated) as the “MBW” complex (Kodama et al. 2018; Koes et al. 2005; Winkel-Shirley 2001; Zhu et al. 2015) to activate the expression of flavonoid/anthocyanin structural genes (LaFountain and Yuan 2021). In poplar, MYB117, MYB134, and MYB115, together with bHLH131, induce the accumulation of anthocyanins by active the expression of DFR or other structural genes (James et al. 2017; Ma et al. 2021; Yoshida et al. 2015); in addition, PtrMYB57 interacted with bHLH131 and PtrTTG1 to form the “MBW” complex and negatively regulated the biosynthesis of both anthocyanins and proanthocyanidins in poplar (Wan et al. 2017). In addition, the accumulation of anthocyanins and proanthocyanidins are regulated by other R2R3-MYB, such as MYB182 (Yoshida et al. 2015), PtoMYB170 (Xu et al. 2017), PtrMYB012 (Kim et al. 2018), MYB6 (Wang et al. 2019), MYB118 (Wang et al. 2020a), MYB165 and MYB192 (Ma et al. 2018). In this study, the transcriptomic analysis showed that the 9 genes (3 MYB113, 3 bHLH, and 3 WD40 domain protein genes) upregulated (Supplementary file 9), and RT-qPCR assays illustrated that 5 of them (1 MYB113, 3 bHLH, and 1 WD40 domain protein genes) upregulated in poplar sunlight-exposed ARs (Figure 6); however, these 3 bHLH (homology with *At*bHLH38 or *At*bHLH16) did not associate with the biosynthesis of flavonoid/anthocyanin in plants. Therefore, this result suggested that sunlight exposure induces the accumulation of flavonols, flavones, and anthocyanins through the activity of transcriptional factors MYB113.

Previous research also showed that the biosynthesis of flavonoids/anthocyanins is regulated by other repressors, such as miR156-SPL and COP1-HY5 module (LaFountain and Yuan 2021). By destabilizing the “MBW” complex, SPLs negatively regulate the biosynthesis of flavonoids/anthocyanins in plants (LaFountain and Yuan 2021). Because some SPLs also are the targets of miR156, then this microRNA can positively control the level of flavonoids/anthocyanins in citrus and switchgrass (Cai et al. 2022; Feng et al. 2022) and poplar (Wang et al. 2020b). Our recent research also illustrated that miR156-SPL modules are involved in the biosynthesis of flavonoids or the development of mosaic symptoms in poplar-BCMV interaction (Wang et al. 2023). Specifically, the expression of miR156 and a series of flavonoid/anthocyanin biosynthesis-related genes (including PAL, CHS, CHI, F3H, DFR, ANS, LAR, and ANR) were upregulated and the expression of four SPL genes (such as gene PAYG024925) were downregulated, however, none R2R3-MYB gene or “MBW” complex significant changed in the poplar mosaic leaves (Wang et al. 2023). Interestingly, the same SPL gene (PAYG024925), which was identified in poplar mosaic leaves, was also downregulated in the present poplar ARs (Supplementary file 9), suggesting sunlight exposure increasing the accumulation of flavonoids metabolites at the post-transcriptional level, through miR156-SPL module which increases the stability of “MBW” complex. In *Arabidopsis*, light signal transduction transcriptional factor HY5 positively regulates anthocyanin biosynthesis through different pathways: directly activates structural genes by binding their promoters (Shin et al. 2013), activates miR858a, which targets the anthocyanin repressor MYBL2 (Wang et al. 2016); and directly binds the promoter of MYBL2 and represses its expression through histone modifications (Wang et al. 2016). As a central switch of light-responsive genes, COP1 represses anthocyanin biosynthesis through destabilizing HY5 and the “MBW” complex (Gangappa and Botto 2016). Then, the HY5 gene (PAYG035590) positively regulates the biosynthesis of anthocyanins, however, the role of the upregulation of the COP1 gene (PAYG018730) in anthocyanins biosynthesis in poplar sunlight-exposed ARs still needs more investigations.

Finally, this study suggests a possible method for the research of flavonoid/anthocyanin biosynthesis. For example, combined with metabolite analysis, the influence of the light spectrum, light intensity, exposure time, etc., on the biosynthesis of flavonoid/anthocyanin; the roles of genes encoding transcription factors, suppressors involved in flavonoids/anthocyanins biosynthesis pathway will be easily conducted using our poplar AR-light induction system.

## Conclusions

In conclusion, this study reports a novel pathophysiological phenomenon in poplar: phloem girdling inoculation with a fungal canker pathogen induces red AR formation on stems in sunlight. Integrated analysis of chemical, flavonoid, and anthocyanin metabolomic analysis revealed that sunlight increases the accumulation of flavonols, flavones, and anthocyanins, while does not change the content of proanthocyanidins. Results also showed that anthocyanins play key roles in the color of poplar ARs, and cyanidin-3-O-glucoside is the colorant of red poplar ARs induced by canker pathogens. Metabolomic and transcriptomic analysis suggested sunlight redirected the metabolic flux of flavonoid biosynthesis in poplar ARs: 1) triggered a rapid anthocyanins biosynthesis, catalyzed by the upregulated BZ1 gene; 2) redirected the metabolic flux from the flavonoid biosynthesis pathway to flavonols branching pathway (by the upregulated FLS gene) and 3) to flavones branching pathway. Results showed that structural genes ANS, BZ1, and FLS play committed roles in the biosynthesis of flavonols, flavones, and anthocyanins, while BZ1 plays a crucial role in the coloration of poplar ARs. Results suggested that sunlight increases the accumulation of flavonoid metabolites through the activity of transcriptional factor MYB113, results also implied that module miR156-SPL and COP1-HY5 involve in the regulation of flavonoid/anthocyanin biosynthesis in poplar ARs. Considering the diverse and important roles of flavonoids on the development of plants, this study not only helps to reveal the coloration of poplar ARs but also benefits the investigation of plant root development, light acclimation, photoprotectivity, responses to pathogens and insects. In addition, this study also provides a possible method for research on flavonoid/anthocyanin biosynthesis.

## Data availability statement

Transcriptomic resources described in this paper are available in NCBI as raw RNA-Seq under the BioProject Accession PRJNA932604. The original contributions presented in this study are included in the article/supplementary material, further inquiries can be directed to the corresponding authors.

## Acknowledgments

The work was supported with funds to Jiaping Zhao from the Central Public-interest Scientific Institution Basal Research Fund of State Key Laboratory of Tree Genetics and Breeding (Grant No. CAFYBB2020ZY001-2), and the National Natural Science Foundation of China (Grant No. 32171776).

## Author contribution

JZ conceived and designed the original research plans. XS supervised the experiments. ML performed most of the experiments. JL, WS, LW, ZL, and SZ provided technical assistance to ML. ML and YF analyzed the experimental data. ML, JZ and HL interpreted the results and wrote the article. All authors have read and approved the manuscript.

## Publisher’s note

All claims expressed in this article are solely those of the authors and do not necessarily represent those of their affiliated organizations, or those of the publisher, the editors, and these viewers. Any product that may be evaluated in this article, or claim that may be made by its manufacturer, is not guaranteed or endorsed by the publisher.

## Supplementary Files

Supplementary file 1. Relative quantification of flavonoid metabloites in poplar ARs.

Supplementary file 2. Quantification of anthocyanin metabolites in poplar ARs.

Supplementary file 3. Validition of the contents of anthocyanin metabolites in poplar Ars

Supplementary file 4. Primer sequences used in this study.

Supplementary file 5. The number and length of fibrous root of pathogen-induced ARs.

Supplementary file 6. Pigment determination in poplar ARs.

Supplementary file 7. KEGG pathway enrichment analysis revealed the differentially expressed genes (DEGs) and differentially accumulated metabolites in flavonoid and anthocyanin biosynthesis pathways enriched in poplar ARs. Heatmap of flavonoids accumulation (A), heatmap of anthocyanins accumulation (B), KEGG pathway enrichment analysis of flavonoids metabolites (C), and anthocyanins metabolites (D) of poplar ARs.

Supplementary file 8. Differentially expressed genes identified in poplar ARs induced by canker pathogen.

Supplementary file 9. RT-qPCR validation of expression of genes involved in the flavonoid/anthocyanin biosynthesis pathway.

Supplementary file 10. The color change of the crude extract of poplar sunlight-exposed ARs under different pH conditions.

