## Supplementary file 5 for "Why the adventitious roots of poplar are so colorful: RNAseq and metabolomic analysis reveal flavonols, flavones, and anthocyanins accumulation in canker pathogens-induced adventitious roots in poplar"

**Supplementary file 5. The number and length of fibrous roots of pathogen-induced ARs (n ≥ 14).**

| **Treatment** | **Number of fibrous roots** | **Length of fibrous roots** |
| --- | --- | --- |
| **Bdo** | 9.857 ±1.351 | 1.335 ± 0.262 cm |
| **S-Bdo** | 11.400 ± 1.724 | 1.198 ± 0.238 cm |
