## Supplementary file 6 for "Why the adventitious roots of poplar are so colorful: RNAseq and metabolomic analysis reveal flavonols, flavones, and anthocyanins accumulation in canker pathogens-induced adventitious roots in poplar"

**Supplementary file 6. Pigment determination in poplar ARs.**

|  | Pigments | Sunlight (mg·g^-1^) | Shading (mg·g^-1^) |
| --- | --- | --- | --- |
| Experiment 1 | Anthocyanidins | 5.91 ± 0.62 | 0.48 ± 0.08 |
|  | Flavonoids | 9.03 ± 1.38 | 5.64 ± 1.05 |
|  | Carotenoids | 0.24 ± 0.03 | 0.10 ± 0.01 |
|  | Procyanidins | 24.18 ± 4.44 | 23.37 ± 2.26 |
| Experiment 2 | Anthocyanidins | 5.98 ± 0.64 | 1.84± 0.35 |
|  | Flavonoids | 9.85± 0.92 | 6.06± 1.20 |
|  | Carotenoids | 0.11± 0.01 | 0.08± 0.02 |
