## Supplementary file 7 for "Why the adventitious roots of poplar are so colorful: RNAseq and metabolomic analysis reveal flavonols, flavones, and anthocyanins accumulation in canker pathogens-induced adventitious roots in poplar"

**Supplementary file 7. KEGG pathway enrichment analysis revealed the differentially expressed genes (DEGs) and differentially accumulated metabolites in flavonoid and anthocyanin biosynthesis pathways enriched in poplar ARs.** Heatmap of flavonoids accumulation (A), heatmap of anthocyanins accumulation (B), KEGG pathway enrichment analysis of flavonoids metabolites (C), and anthocyanins metabolites (D) of poplar ARs.

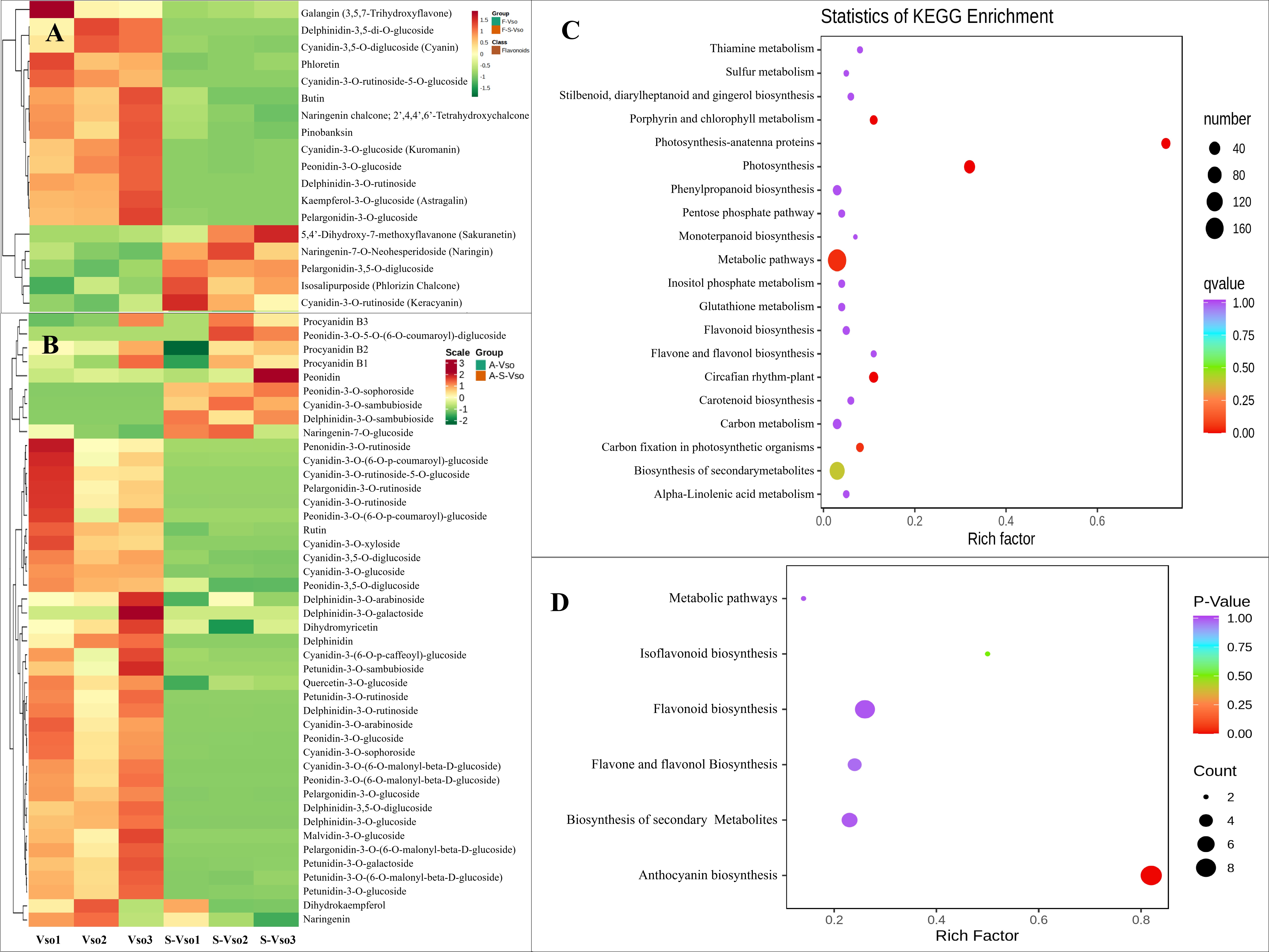
