## Supplementary file 10 for "Why the adventitious roots of poplar are so colorful: RNAseq and metabolomic analysis reveal flavonols, flavones, and anthocyanins accumulation in canker pathogens-induced adventitious roots in poplar"

**Supplementary file 10. The color change of the crude extract of poplar sunlight-exposed ARs under different pH conditions.**

**[Methods and materials]** Plant materials used in this study are *P. alba* var. *pyramidalis* saplings cultivated in pots containing a growth substrate of peat and perlite (v:v = 4:1) in the experimental field of the Chinese Academy of Forestry (Beijing, China).Canker pathogen used in this study is*B. dothidea*strain CZA, grown on 2.0% PDA at 30 ℃ in the dark for 7 days. Inoculation method is same as the text. After pathogen inoculation, the girdling sites were wrapped with polyethylene (PE) film to maintain moisture. After the formation of poplar AR, 3.0 g AR sample (formed in sunlight or shaded conditions) was heated in 3 mL deionized water at 100°C for 5 min for crude pigment extraction. The supernatant (crude pigment solution) was evenly divided into 6 tubes. Then, 0.5 and 1.0 μL of 1 MHCl solution and 0.5, 1.0, and 5.0 μL of 1 M NaOH solution were added to the pigment extract in five tubes; the sixth tube, to which no reagents were added, was used as the control. The color change under different pH conditions was observed and photographed.

**[Results]** To validate the color stability of anthocyanins under different pH conditions, the crude extract of red poplar ARs was acidified and alkalified by HCl and NaOH, respectively. As shown in Figure S1, the color of crude pigments extracted from red ARs were pink; and the acidified pigments (~ pH 2.7 and pH 3.0, respectively) were coral red; the alkaline pigments (pH 11.0, 11.3 and 12.0, respectively) were purple or deep purple. In addition, the crude extract of poplar shadedARs was milky white, and no color change was observed in the acidic and alkaline solutions (Figure 1).This result suggests that anthocyanins are colorants of poplar ARs and that no anthocyaninmetabolitesare produced under shadedconditions. Based on the above results,metabolomic analysis of anthocyanins and flavonoids in poplar ARs wasconducted in this study.

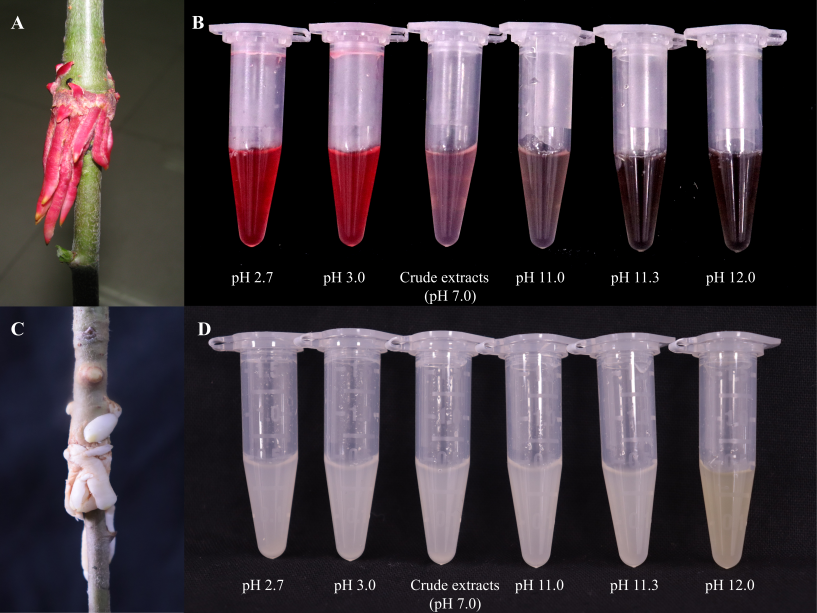

Figure S1. The color change of the crude extract of poplar sunlight-exposed ARs under different pH conditions. Poplar ARs induced by canker *Botryosphaeria dothidea* in under light conditions (A); the color of the pigment extract varied under pH conditions (B). ARs induced under shaded conditions (C); the color of the extracted pigments did not vary under pH conditions (D).
